# SPrOUT: A computational and targeted sequencing approach for mixed plant DNA identification with Angiosperms353

**DOI:** 10.64898/2026.02.20.707031

**Authors:** Nan Hu, Madison Bullock, Chris Jackson, Courtney Miller, Elizabeth Sage Hunter, Charles Huff, Yanni Chen, Sara M. Handy, Matthew G. Johnson

## Abstract

**Premise:** The identification of plant species from mixed samples is crucial in various fields, including ecological surveys, conservation efforts, and food and dietary supplement safety. Traditional methods face potential challenges due to the high costs of DNA sequencing, inefficiencies in computational workflows, and incomplete sequence databases.

**Methods and Results:** This study introduces a novel approach using the Angiosperms353 target sequencing kit for efficient taxonomic identification of angiosperm DNA in mixed samples. Our method assembles short pair-end reads for each mixed sample. Using gene sets of Angiosperms353 from 871 species, we apply phylogenetic inference to categorize the variance in phylogenetic distance across genes to identify the presence of taxa in mixed plant samples. The pipeline reaches 98.1 to 99.6% accuracy, 92.9 to 100% precision for identifying unknown taxa in in-silico mixes, and 90.7% accuracy and 98.0% precision for mock supplement mixtures. We explored the parameter cutoffs of the pipeline to offer an empirical range for different applications.

**Conclusions:** The Angiosperms353 and HybPiper assembly proved effective in sorting mixed plant DNA samples. Our method offers a framework for scientific and practical applications in plant species identification in both single and mixed samples.

## INTRODUCTION

From habitat shifts to the spread of invasive species, global biodiversity has faced escalating threats across multiple scales over recent decades (Newbold, 2018). These shifts underscore an urgent need for robust methods in species identification, conservation monitoring, and ecological assessment to effectively document and mitigate biodiversity loss. Accurate taxonomic identification is essential not only for advancing scientific understanding and conserving biodiversity but also for safeguarding public health and maintaining secure food supply chains (Bortolus, 2008). Despite the growing importance of such identification, the absence of a reliable, universal classification pipeline for plant species identification presents a significant bottleneck.

Traditional plant species identification has long relied on either morphological traits or single-gene DNA barcoding, each presenting unavoidable limitations. Morphological identification, while useful, is labor-intensive, requires specialized expertise, and struggles to identify plants when samples are fragmented, degraded, or lacking distinctive morphological traits (Crisci et al., 2020). DNA barcoding, which emerged as a popular method in plant systematics, typically targets plastid genes (such as *rbcL* and *matK*) and ribosomal DNA sequences, as these loci are universally present in most land plants (Fazekas et al., 2008; Hollingsworth, Graham and Little, 2011; Newbold, 2018). Plastid and ribosomal sequences offer practical advantages for single-species identification: they are relatively easy to sequence and have high copy numbers, facilitating identification from small DNA samples. However, although plastid and ribosomal DNA provide reliable markers for some groups, they often lack sufficient resolution to distinguish closely related species, resulting in ambiguous or even incorrect identifications in certain taxonomic groups (Yu et al., 2022). For example, plastid genes can be highly conserved within genera or families, limiting their utility in fine-scale taxonomic resolution.

These issues are further complicated in metabarcoding applications, where samples contain mixed DNA from multiple species. Metabarcoding has achieved notable success in identifying microbial and animal species (Norgaard et al., 2021; Antil et al., 2023; Schutte, Stuben and Astrin, 2023; Hebert et al., 2025). However, comparable advances in plant metabarcoding are constrained by several challenges (Chac and Thinh, 2023) and existing tools consistently underperform with complex angiosperm mixtures (Piper et al., 2019; Liu et al., 2020; Schutte, Stuben and Astrin, 2023). Reliance on plastid markers can lead to spurious results due to the high copy number and limited variability of these organellar genomes. Additionally, methodological challenges such as primer bias, short amplicons, and degraded DNA, can lead to misidentifications or false negatives of taxa (Bell et al., 2016; Bruno et al., 2019; Arstingstall et al., 2021). The main advantage of barcoding and metabarcoding approaches is that specific chloroplast genes and the ITS regions are abundant in public databases, while whole genome representation of angiosperms remains sparse.

As next-generation sequencing has become cheaper and more prolific, researchers have tried to increase the availability of reference angiosperm genomes. Unfortunately, this progress is limited by the size and complexity of plant genomes, and the general lack of plant nuclear genome references remains a hindrance for the development of new computational approaches (Alsos et al., 2018; Vallin et al., 2025). Reference free approaches, such as genome or chloroplast skimming, while offering more sequence data, often fail when working with processed or chloroplast-poor tissues, and remain expensive when using deeper nuclear-genome sequencing to resolve close relatives (Hunter, Literman and Handy, 2021; Wizenberg et al., 2023).

Recently, nuclear protein-coding genes have emerged as valuable candidates for plant species identification in complex samples. These nuclear markers have the capacity to distinguish closely related species, and capture genetic variation linked to key morphological traits (Zimmer and Wen, 2015; Pezzini et al., 2023). However, their utility has been constrained by several factors: (1) the lack of a universally conserved gene set across major plant groups; (2) limited availability of reference databases; (3) underdeveloped computational tools for data analysis; and (4) the comparatively high cost and complexity of sequencing nuclear genes relative to plastid genes (Chac and Thinh, 2023). Fortunately, target capture sequencing methods can ameliorate many of these challenges.

Angiosperms353 is an ideal platform to explore plant identification studies. The universal design of Angiosperms353 is probe sequences targeting 353 nuclear protein coding genes found at low copy in flowering plants (Johnson et al., 2019). Since its development, Angiosperms353 has demonstrated remarkable robustness and adaptability in plant systematics, validated by hundreds of studies that highlight its effectiveness across diverse plant lineages and phylogenetic scales from orders to populations (Lee et al., 2021; Maurin et al., 2021; Perez-Escobar et al., 2021; Siniscalchi et al., 2021; Slimp et al., 2021; Thomas et al., 2021a; Thomas et al., 2021b; Zuntini et al., 2021). Although many of these studies generate sequences via the targeted sequencing approach, the Angiosperms353 loci can also be found by mining data from transcriptomes and genomes, including some gymnosperms (Baker et al., 2021). The Plant and Fungal Tree of Life (PAFTOL) project used Angiosperms353 data from over 10,000 species to construct one of the largest flowering plant phylogenies to date (Zuntini et al., 2024). The maintenance of Angiosperms353 data on publicly available websites provides a valuable database against which unknown sequences may be compared. However, the potential for using Angiosperms353 as a plant identification tool in metagenomic applications remains largely unexplored.

Compared to conventional approaches, using nuclear genes from a target capture dataset with a phylogenetic-distance-based approach has clear advantages in plant species identification. Targeted sequencing methods overcome the biases of single-gene PCR-based pipelines (China Plant BOL Group et al., 2011), can be used to assemble sequences even with fragmented DNA (Brewer et al., 2022) and can generate data for mixed DNA cost-effectively (Hale et al., 2020; Bullock et al., 2025). and integrate cumulative phylogenetic signals across the nuclear genome rather than relying on a few marker genes. However, most bioinformatics workflows for metagenomics are developed to reference only a few genes (Schloss et al., 2009; Bolyen et al., 2019). By targeting hundreds of markers across the nuclear genome, targeted sequencing has the potential to increase the potential for identification by deriving phylogenetic distances against a comprehensive reference phylogeny.

To explore this potential, we introduce SPrOUT (Species PRediction Of Unknown Taxa) as a comprehensive solution to single or mixed plant DNA identification, including guidance for data generation and computational pipelines for taxonomic prediction. This approach is designed based on phylogenetic distance methods to overcome current barriers of the identification process, reducing associated costs, and enhancing taxonomic resolution in complex plant samples. Through this methodology, we contribute a valuable tool for conservation biology, ecological monitoring, environmental biology, and food safety, thereby advancing plant metabarcoding as a precise and accessible means for species identification in mixed-sample contexts.

## METHODS AND RESULTS

### Overview

SPrOUT is a Linux-based Python workflow designed to predict plant species from target capture sequencing data, applicable to both single and mixed samples (Figure 1). Our pipeline has 4 major parts: Data processing, Target assembly, Phylogenetic Inference, and Prediction. Users can start with either raw tissue samples (to build a target sequencing library) or preexisting short-read data for target gene assembly. The current implementation includes two reference sets in different taxonomical hierarchies: 106 taxa for order level prediction and 298 for family level prediction, which both subset from precisely curated Angiosperms353 genes from 871 plant sequencing arrays. The pipeline outputs the closest taxonomic assignment of the input data based on cumulative phylogenetic signals from the target region data.

**Figure 1.**
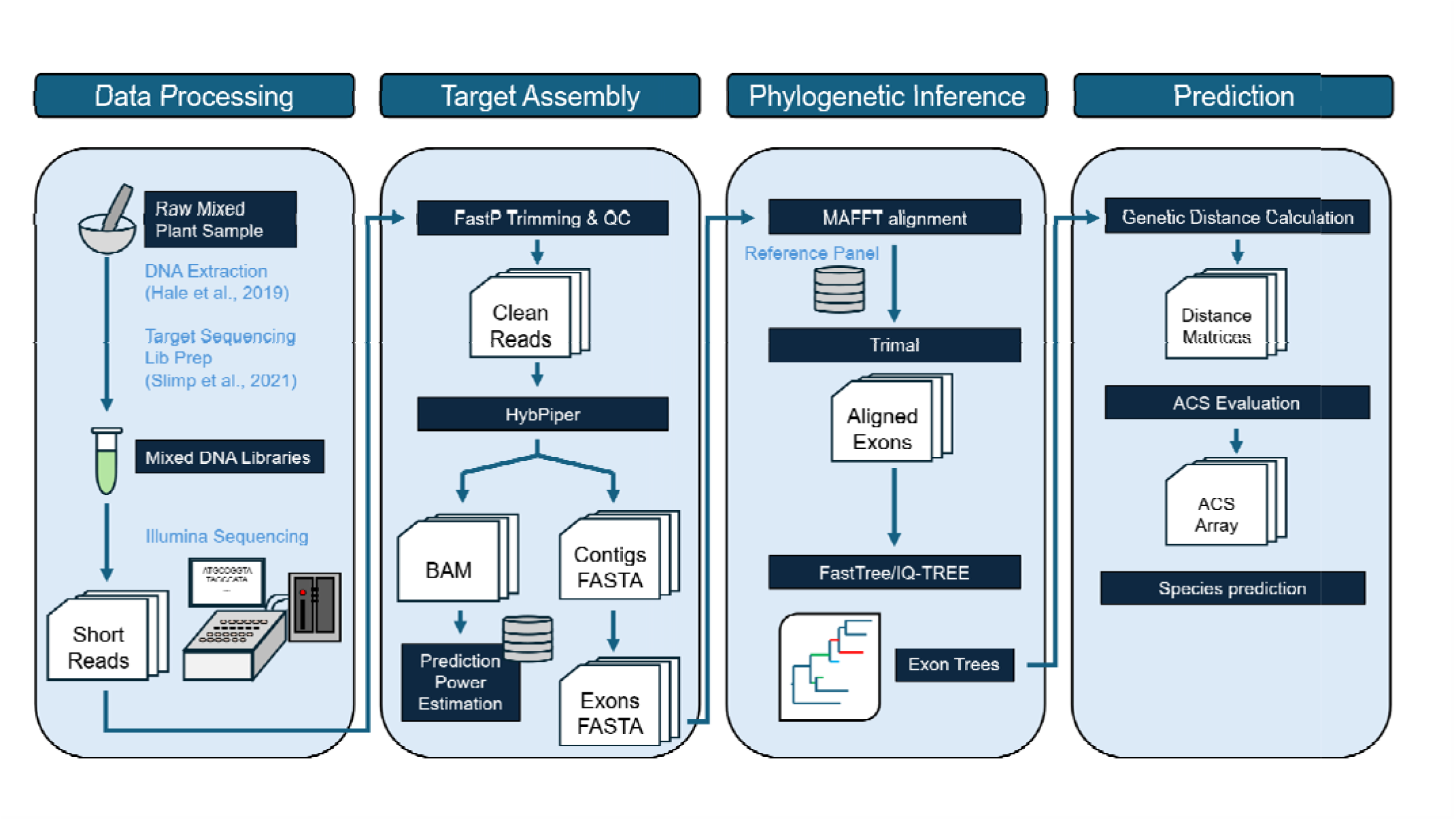
Workflow of the pipeline. The four procedures of the pipeline include: data processing, target assembly, phylogenetic inference, and prediction.

### Reads assembly and exon prediction

We began by processing the input raw sequencing reads for trimming and quality control using Fastp (Chen, 2023), removing sequencing adapters and low-quality reads (Q score < 25). To recover the Angiosperms353 genes, we used HybPiper v2.2.0 (Johnson et al., 2016), with post-quality control reads as input. We specified the target file “mega353” for the HybPiper assembler, available at https://github.com/chrisjackson-pellicle/NewTargets. HybPiper maps reads to reference target sequences, and then performs de-novo assemblies of the mapped reads. The “exonerate” function (v2.4.0) within HybPiper predicts exon boundaries for each assembled contig (Slater and Birney, 2005). The exon boundary data are then used within customized python scripts to generate exon sequence files for each targeted gene. Default settings require a minimum of 80% overlap for recognizing identical exons; this argument is adjustable in the pipeline

### Phylogenetic inference

We align exons to genes from the reference panel, obtainable through whole-genome assembly via BLASTX v2.15.0 (Camacho et al., 2009), or from other targeted sequencing projects (Goodstein et al., 2012; One Thousand Plant Transcriptomes, 2019). The latter can be converted into FASTA sequences using the “hybpiper retrieve_sequences” function after assembly. We conduct the sequence alignment using MAFFT v7.526in “—addfragments” option (Katoh and Frith, 2012). After alignment, we trimmed the aligned FASTA files with trimAl v1.5.0 (Capella-Gutierrez, Silla-Martinez and Gabaldon, 2009) using the “-gt 0.5” setting to eliminate unrelated sequences. For tree inference, we used three approaches of increasing complexity: FastTree v2.1.11 using the GTR+Gamma model (Price, Dehal and Arkin, 2009), IQ-TREE v2.3.6 with a fixed evolution model (Minh et al., 2020), and IQ-TREE with model finder (MFP) (Kalyaanamoorthy et al., 2017). Each alignment produced an exon tree with associated node support values.

### Genetic distance and species relation prediction

We processed the exon trees to calculate pairwise genetic distances under a specified evolutionary model. The re-rooting group can be either user-selected outgroups or by midpoint. To eliminate the misaligned exons, we excluded lineages with abnormally long branch lengths based on Z-test results, taking node support into account (see Figure S1). We then summarized pairwise distances between unknown exons and the reference panel across all exon trees to compute an Adjusted Cumulative Similarity (ACS) (See Table 1 for Glossaries). We based our species predictions on a Z-score percentile, assuming ACS distribution follows a normal curve. A higher ACS score indicates a higher likelihood of close relation to the reference species.

**Table 1.** Glossary of terms.

| Terms | Description | Equation |
| --- | --- | --- |
| TPR | True Positive Rate, recall, sensitivity, probability of detection, hit rate, power. A measurement of the proportion of predicted true positive events among all the actual positive events. This evaluates the ability of identifying correct existing components in the mix by the pipeline | $TPR = \frac{True\ Positive}{Actual\ Positive}$ |
| TNR | True Negative Rate, specificity, selectivity. A measurement of the proportion of the predicted true negative events among all the actual negative events. | $TNR = \frac{True\ Negative}{Actual\ Negative}$ |
|  | This evaluates the ability to reject non-existing species in the mix. |  |
| PPV | Positive Prediction Value, precision. A measurement of the proportion of true positive events among all the predicted positive events. This represents the degree of how much we can trust the positive predictions from the pipeline.<br>Mathematically, PPV is equal to the probability of predicted species is the true existing species in the mix. | $PPV = \frac{True\ Positive}{True\ Positive + False\ Positive}$ |
| NPV | Negative Prediction Value. A measurement | $NPV = \frac{True\ Negative}{True\ Negative + False\ Negative}$ |
|  | of the proportion of true negative events among all the predicted negative events. This represents the degree of the model to identify true negative events among all negative events. We use this evaluation in this pipeline to adjust the amount of taxonomy groups involved in hierarchical prediction. |  |
| ACC | Accuracy. A measurement of the proportion of both true positive and negative predictions in total actual condition. This is an overall indicator of the performance of the | $ACC = \frac{True\ Positive + True\ Negative}{Actual\ Positive + Actual\ Negative}$ |
|  | pipeline. |  |
| ACS | Adjusted Cumulative Similarity. A transformed evaluation measuring how closely the testing samples are related to known references. This takes the phylogenetic distance from the tree to calculate the pairwise similarity for each exon tree. Then, the result is filtered by a decision tree (See Figure S1) and summarized from all exon trees. This number serves as the final determinator of the species prediction., | $ACS = \sum \frac{1}{1 + Genetic\ Distance}$ |
| Z-score | Assuming the data distribution is normal, Z-score represents how | $Z = \frac{ACS - Mean(ACS)}{Standard\ Deviation}$ |
|  | many standard deviations an observation is from the mean. High Z-score means the single data is an outlier in the population, which in our case, the candidate of relative species to the certain reference. |  |

### Species prediction evaluations from in-silico libraries

To examine the predictability of our pipeline, we performed tests with both the in-silico mixes and known mixed samples. For the in-silico tests, we selected 30 species from Kew Garden Tree of Life database (Zuntini et al., 2024) (Table S1). Reads for all 30 species reads were generated via target capture arrays (with PAFTOL or equivalent methods marked on Tree of Life Explorer). Our selection aimed to cover major angiosperm clades and maximize lineage coverage. We created three different test datasets from the 30 species. (1) We generated 100 samples of artificial mixed reads, each consisting of mixed reads from three, six, or ten species in equal proportion (Table S2). (2) We produced 36 samples with skewed composition, varying both the species composition and read counts (Table S3), to test the stability of the pipeline in unevenly mixed scenarios where dominant samples might interfere and mask the minor components in the mix. (3) We prepared 36 low-yield samples by subsampling at different reads mapping recovery levels (Table S4), estimating recovery rates from BAM output of the HybPiper using ‘samtools flagstat’ v1.20 (Li et al., 2009), and generating reduced read files with seqkit v2.82 (Shen, Sipos and Zhao, 2024). For each experimental scenario, we created replicates with different random seeds, as documented in Table S3 and Table S4.

#### Single species samples

To test whether the pipeline could accurately identify the 30 test species in single-species read sets, we validated the pipeline on single-species libraries and correctly identified all 30 test species at both order and family level by selecting the species with the highest ACS. Among 30 testing species, 29 (96.7%) achieved ACS values at least 7 or more standard deviations (SD) above the mean, categorizing them as significant outliers (p < 10^-12^), while only one showed a lower ACS value of 4.03 (Malvales, p < 10^-5^). These results demonstrate the high reliability of the pipeline for single-species identification. Because our test sets included diverse taxonomic groups, our single species analyses also established a reliable baseline for evaluating subsequent mixed-sample predictions.

#### Mixed samples

For 100 artificial mixes composed from the PAFTOL database, we evaluated our predictions based on multiple factors from the confusion matrix. The True Positive Rate (TPR, recall, sensitivity), True Negative Rate (TNR, specificity), Positive Prediction Value (PPV, precision), and the Accuracy (ACC) were calculated based on the statistical results from the predicted closed related species and the actual closed related species in each sample combinations (See Table 1 for Glossaries). Our glossary aligns standard concepts in machine learning with our statistical results to reduce misleading inferences.

We performed multiple tests to determine a practical best range of threshold for the Z-score of ACS to consider a significant positive prediction in all samples (Table 2; Table S5) (Figure 2). We found a generally decreasing trend of TPR when the Z-score of ACS increases, which means more actually present species in the mix are missing from the prediction when we restrict the Z-score standard (Figure 2a). However, TPR – which represents the rate of SPrOUT successfully rejecting non-existing species – was maintained at 90% when the Z-score rose to 1.1 (Mean = 0.900, SD = 0.143). From the Z-score range of -0.1 to 2, the pipeline reached 90% of correctly rejecting negative cases (Mean = 0.948, SD = 0.046, at Z = -0.1) (Figure 2b). We also tracked Positive Predictive Value (PPV), the proportion of true positives (Table 1). With the increase of Z-score, the PPV also increases (Figure 2c). The lowest boundary of Z-score that makes the PPV above 90% is 0.2 (Mean = 0.929, SD = 0.154). The overall evaluation of accuracy (the proportion of species classified correctly) is shown in ACC plot (Figure 2d). When the Z-score is between -0.1 to 2, the ACC is above 90%, with a peak at 0.5 (Mean = 0.996, SD = 0.010). Additionally, these evaluating parameters have distinct performance at different numbers of mixed individuals, especially affecting the variance (Figure S2). We found that evaluations had reduced reliability when the mixture was more complex (more contributing species) compared to simpler compositions. The convincing range of Z-score shrinks into narrower space due to loss or incorrect predictions.

**Figure 2.**
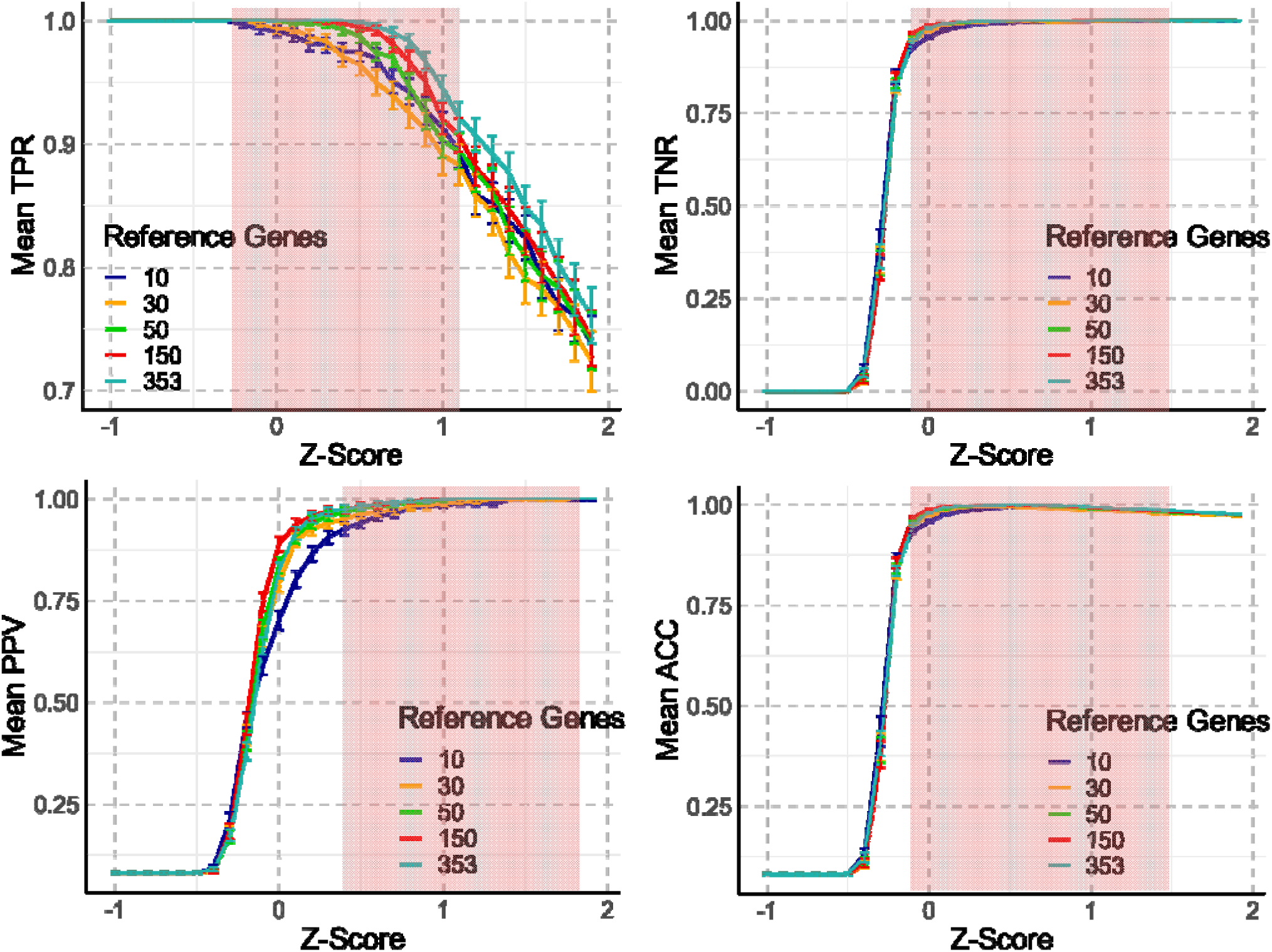
Evaluations of the pipeline using four parameters from the confusion matrix (See Glossary, Table 1). TPR = True Position Rate; TNR =True Negative Rate; PPV = Positive Predictive Value; ACC = Accuracy. Colored lines indicate performance under different number of reference genes for species identifications. Pink shades represent the informative Z-score range where evaluating parameter in each panel is higher than 90%. Error bar is the standard error.

**Table 2.**
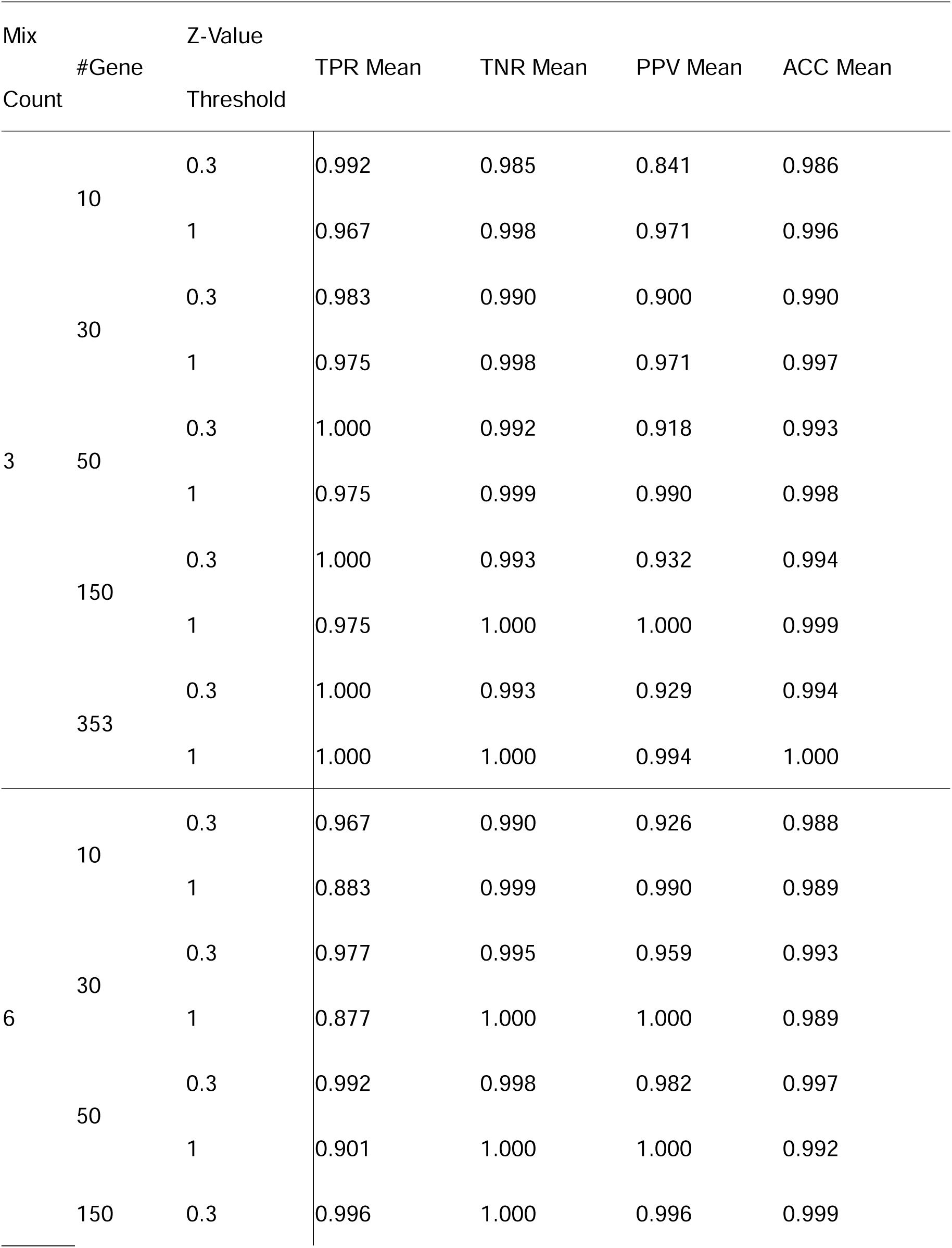
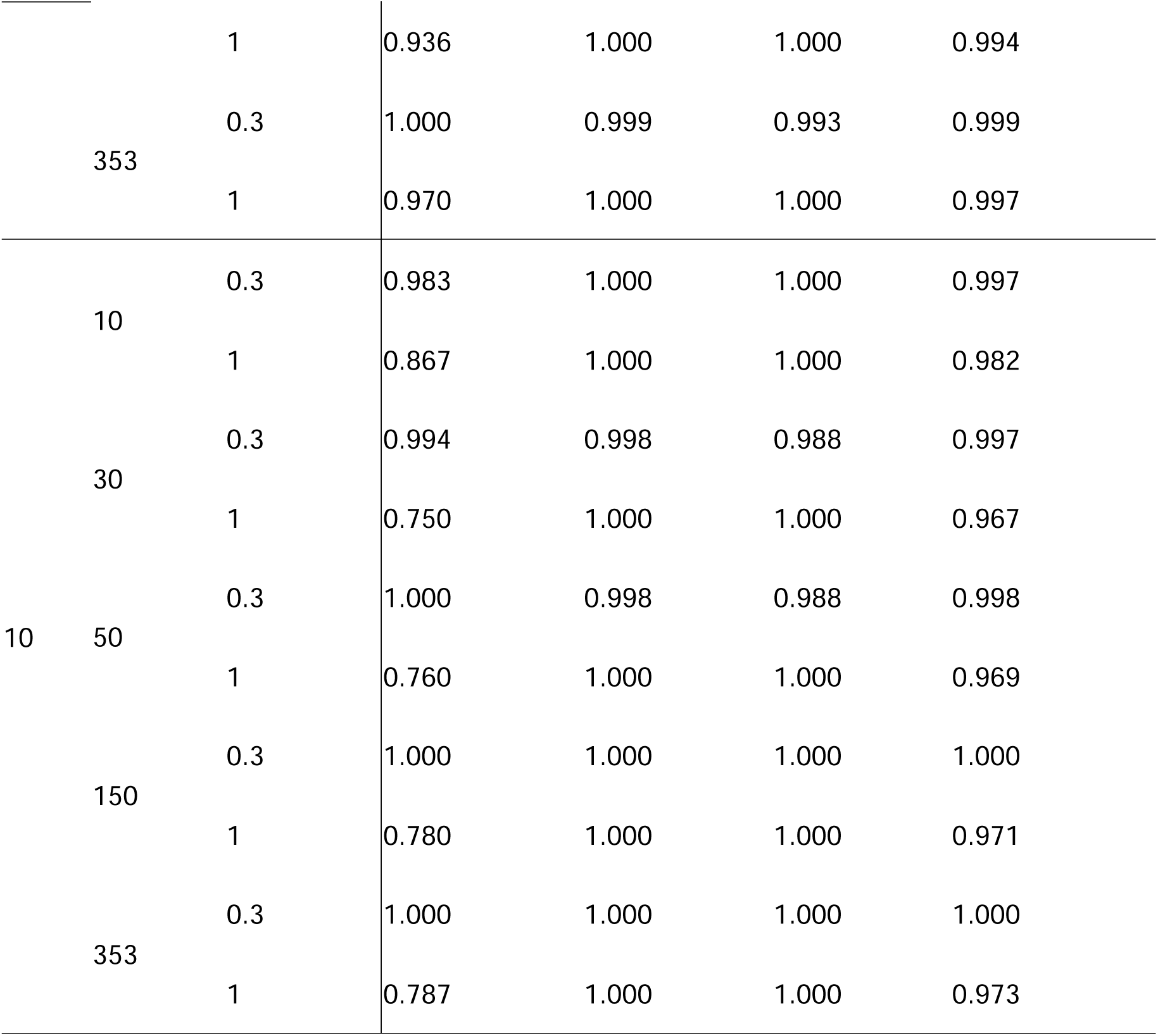
Confusion matrix parameters summary of 100-mix tests.

#### Proportion skewed samples

While the above tests focused on mixed samples with equal contributions, we also performed tests on the predictability with either a skewed proportion or a limited number of input reads. We focused on both the performance from the HybPiper assembler and the final ACS predictions. An unbalanced mixed read input could potentially interfere with the performance of the assembler, which may cause targeted gene loss. In the proportional test, we investigated the impact of a major contributor compared with a minor contributor in mixed samples. We found that a low proportion for one species does not affect correct identification when the total reads number is sufficient (Figure 3a). However, when there are fewer than 20,000 reads mapped to the targets, the lower proportion contributors have significantly worse performance compared to when the same contributor is on its own. (p < 0.01). Table 3 shows that with fewer than 20,000 reads on target, the proportional minor species (Poales) failed to be detected, due to limited genes recovered from the HybPiper; in contrast, while testing samples separately, some assemblies were successfully acquired with only 15,000 reads This test suggests a strong interference in unevenly composed mixed samples when input reads are limited. This also suggests that some source of false negatives may be due to the input reads quantity.

**Figure 3.**
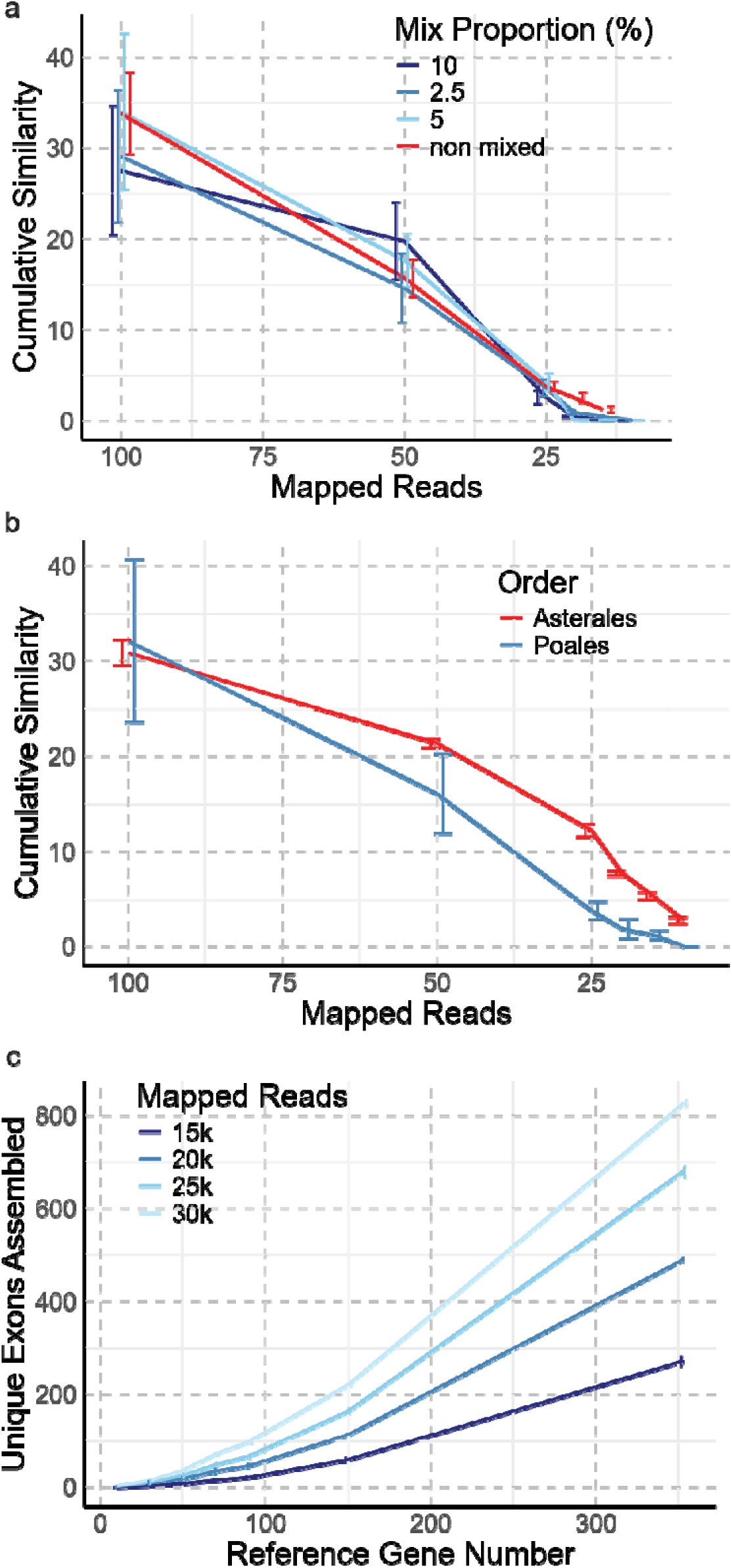
Performance of minor proportion species, low reads count, and reduced reference gene number. (a, b) Cumulative similarity for minor species across decreasing mapped read numbers shown by (a) different mixing proportions (%); (b) Two orders (Asterales and Poales) separately. Cumulative similarities decline when mapped reads count, particularly below 25,000 reads per species, with minor species in the mixed samples failing to recover any assemblies at 20,000 reads or lower (See Table 2 for details). (c) Number of unique exons assembled as a function of reference gene number at varying mapped read depths per sample. Higher mapped read counts consistently yield more exons recovered, especially as the number of reference genes increases. Error bars represent standard error across replicates.

**Table 3.**
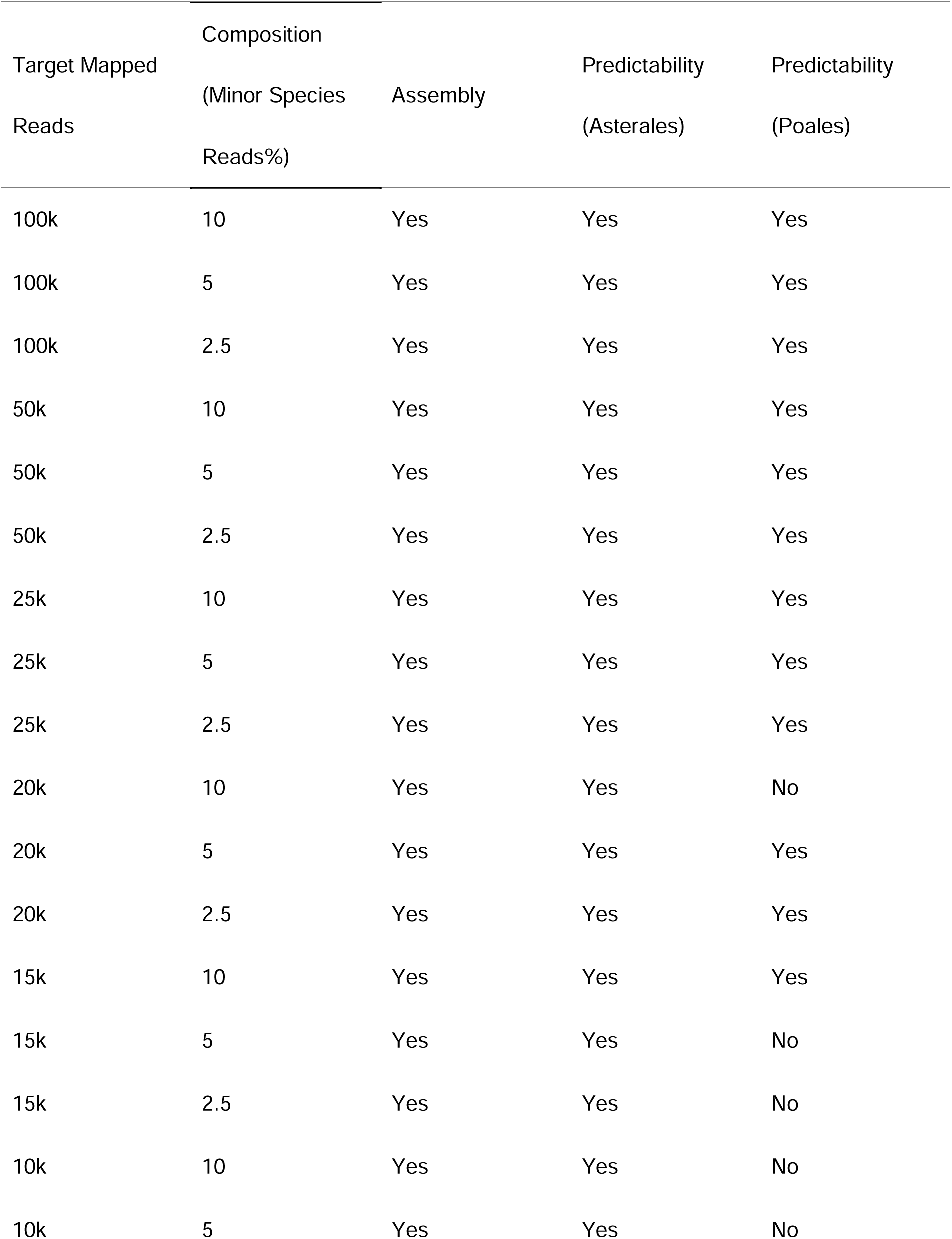
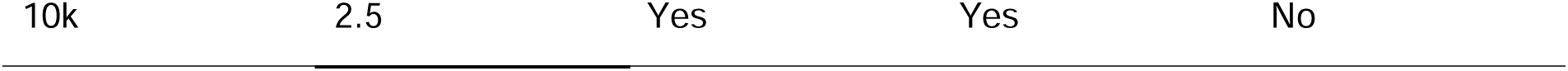
Predictability of pipeline with limited mapped reads and different composition.

#### Low yield samples

We tested the pipeline under a very limited read scenario (10k, 15k, 20k, 25k, 30k reads on target). When the mapped read number was reduced for both an Asterales and a Poales species in this two-species mix, as the reads number decreased, the ACS is linearly dropped for both species (Figure 3b). However, different species have distinct performance on the curve when the number of mapped reads is relatively low (<50,000 mapped reads per species). We also checked the assembled unique exons number under different mapped reads number (Figure 3c). We found that the number of exons that were constructed into a tree drastically decreased when there was very limited input. This factor is strongly correlated with the ACS under different settings of initial mapped reads number (R^2^ = 0.968, p < 0.05).

#### Family level predictions

Our pipeline permits predicting species in a hierarchical way to reduce the computational load and to increase the accuracy. For unknown mixed samples, we can run the pipeline using rough order level references to sort the unknown components into their orders. Then, we select families within the predicted orders to make lower taxonomical predictions. To fulfill the hierarchical method in our pipeline, we further predicted the 60 mixes down to the family level based on the order level results. The composition of these 60 mixes includes within-order mixes, across-order mixes, and 3-order complex mixes (Table S6). The selection of reference families was adjusted by taking cutoffs at 85%, 90%, and 95% empirical NPV level from order level predictions. The family-level prediction heatmaps (Figure 4) visualize the evaluation performance across multiple conditions. The results were analyzed by varying the number of reference genes (10, 30, 50, 150) and adjusting NPV cutoffs (85%, 90%, 95%) from the order-level predictions (See Table 1: NPV for explanation). In Figure 4a, which presents True Positive Rate (TPR) comparisons, the heatmap tiles reveal a consistent trend of higher TPR values concentrated at higher NPV cutoffs (90% and 95%), especially for larger numbers of reference genes (50 and 150). This indicates a marked improvement in prediction sensitivity when stricter cutoffs and larger gene pools are employed. Figure 4b, focusing on Positive Predictive Value (PPV), exhibits a similar gradient. However, higher PPV values are prominently observed in regions corresponding to loosened NPV cutoffs (85%) while larger reference gene pools still contribute to high PPV. This reveals a tradeoff in order level selection – including more orders in the reference data will increase the sensitivity (probability of detection), at the expense of losing specificity (Type I error, probability of a false identification).

**Figure 4.**
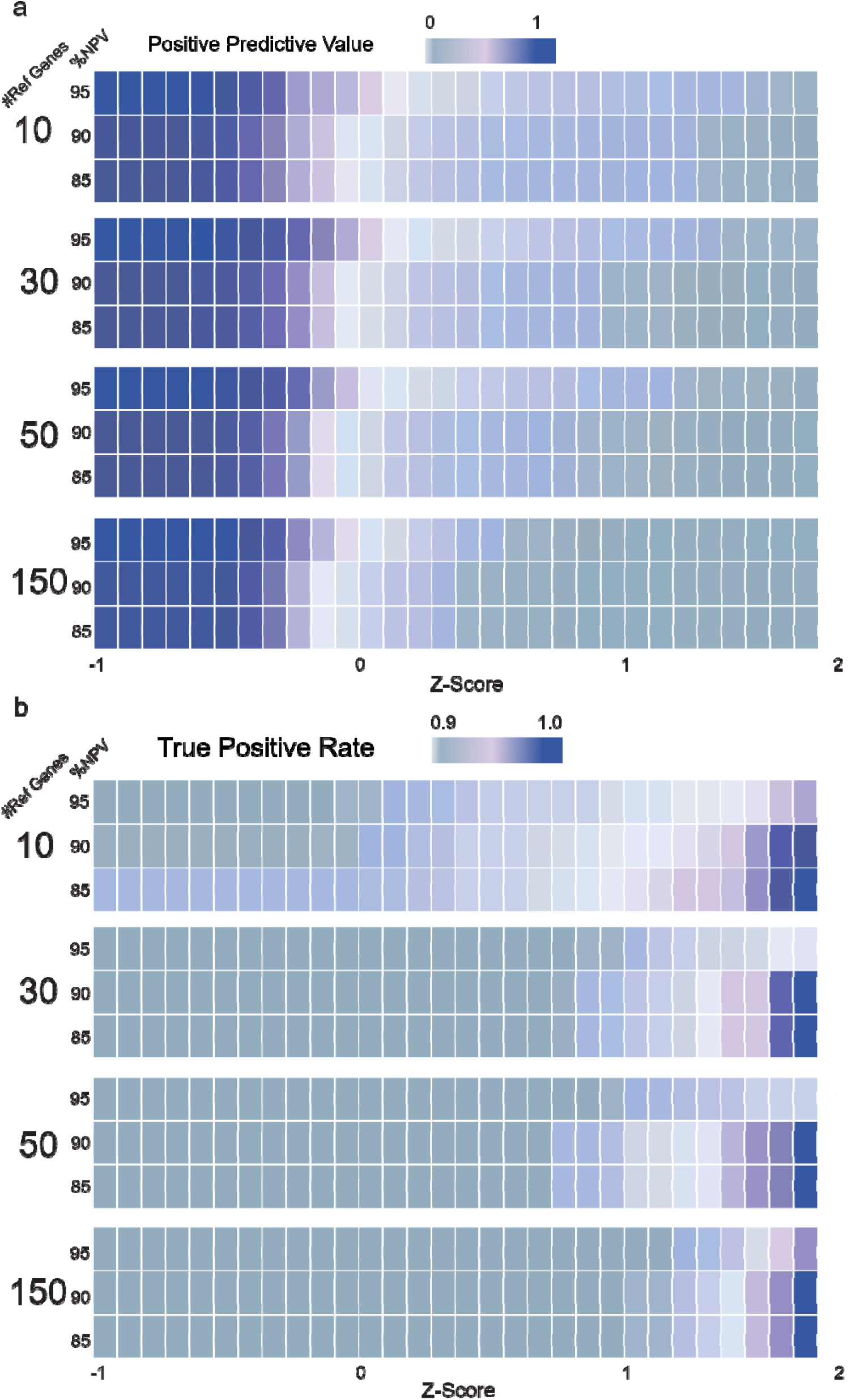
Heatmaps of family level predictions. Colored tiles indicate the value of TPR and PPV evaluated from selected family mixed datasets. For each number of reference genes test, selected orders as candidate pools for families were decided at 85%, 90%, and 95% NPV (Negative Predictive Value) from order level results (a) TPR comparisons among different conditions (b) PPV comparisons among different conditions.

### Species identification from known tissue mixtures

We generated eleven artificial mixed libraries as tests for the real-world samples (Figure 5). We collected plant vouchers from Smithsonian (Jun Wen), Jim Duke, American Herbal Pharmacopeia (AHP), and NCNPR (Table S7). Dry tissues were weighed to ensure the equalized overall proportion of each mixture for every component. They were then ground with an IKA mill (150,000rpm, 30 sec x 2) and turned for an hour in 50mL to homogenize before sub sampling for extraction. Input reads were generated using a modified target sequencing approach (Hale et al., 2020; Slimp et al., 2021). The probe set used for angiosperm species identification is the myBaits Expert Angiosperms-353 (Johnson et al., 2019) (Arbor Biotechnologies, Cambridge, MA, USA). Each mix contained a variety of species, and the expected taxonomy was compared with the predicted taxonomy at both family and order levels (Figure 5a, Table S7). In mixes containing single-genus species like *Brassica* and *Prunus*, the family-level predictions (e.g., Brassicaceae and Rosaceae) were highly consistent with expectations, reflecting the reliability in distinguishing taxa from unrelated groups. Taxonomic predictions at the order level (e.g., Brassicales, Rosales) showed strong agreement with the expected classification. In more complex mixtures, such as Mix1 and Mix5, containing species from multiple families (e.g., Asteraceae, Apiaceae, Zingiberaceae), the predictions reflected a mix of accurate and slightly divergent classifications. A few discrepancies were observed in predictions for species like *Ginkgo biloba* in Mix3, due to reference database limitations (Angiosperms353 data for gymnosperms is sparse, compared with flowering plants). The species identification results underscore the strength in accurately resolving family and order-level taxonomy across diverse mixes, with high accuracy for single-genus and simpler mixtures (Figure 5b). However, challenges remain in differentiating highly divergent or phylogenetically similar taxa within complex samples. Refinements in the reference database and algorithmic sensitivity could further enhance predictive accuracy.

**Figure 5.**
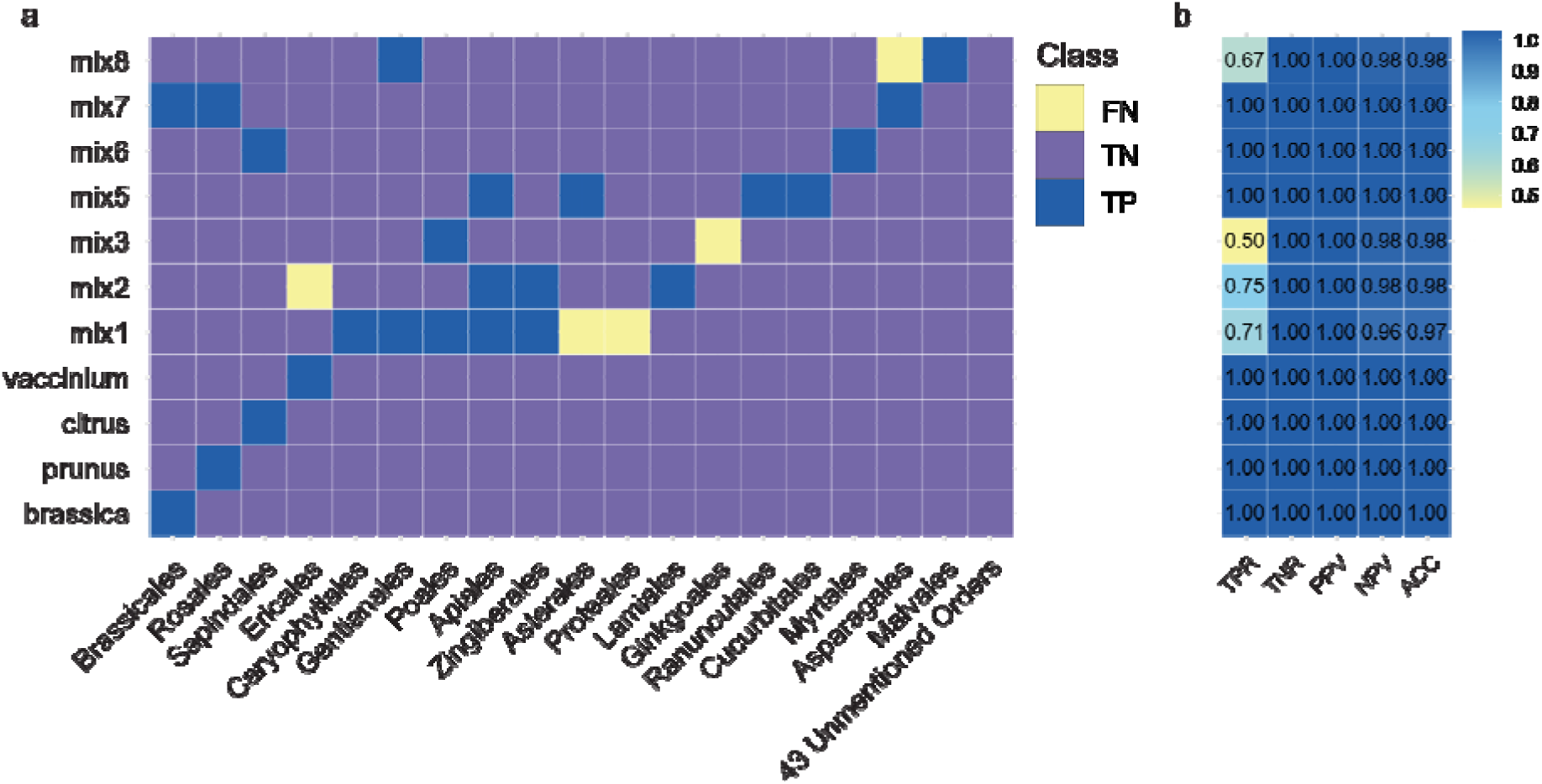
Heatmap and confusion matrix parameters of predictions for real-world mixed samples when Z-score of ACS is between 0 to 1, unless specifically informed. (a) Heatmap of predicting results of 11 real-world mixes. (b) Key confusion matrix parameters in each mixed sample.

### Computing efficiency test

To test the impact of reducing gene number to the pipeline, we performed multiple runs of pipeline with different number of genes (10, 30, 50, 150, and 353). Reduced gene numbers would speed up the phylogeny and distance matrix generation steps drastically, which would in addition reduce the computational intensity of the entire pipeline. All the computational time was estimated under SLURM job queue system in Linux cluster of Texas Tech University High Performance Computer Cluster. Tests were performed with 4Gb input reads, 64Cores AMD EPYC™ 7702 CPU, 512GB RAM with 3.9Gb memory assigned to each CPU core. In different test runs with various input data size and reference choice, the computational time varies from 3 min to hours (Figure 6). The input read size strongly affected the HybPiper efficiency, while tree reconstruction and genetic distance matrix calculation were affected by the number of genes and number of reference species included. The runtime is affected by the input read size and the mapping rate. In Figure S3, we found a linear relationship between mapped read count and the HybPiper runtime. Empirically, when 30 genes were tested using 106 species as references, the total estimated time would be 2-10 min considering the mapped reads ranged in 60-600Mb.

**Figure 6.**
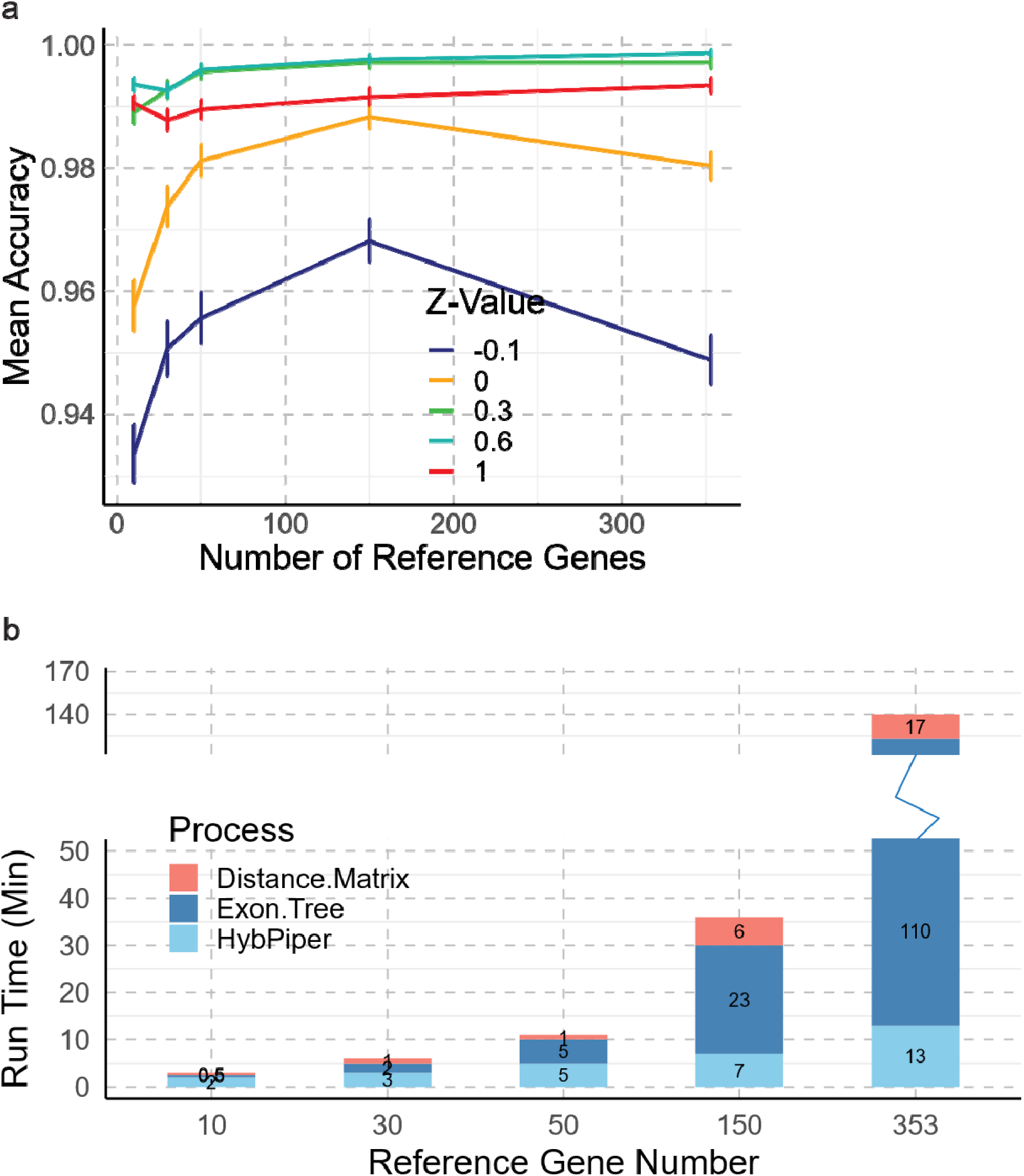
Computational efficiency of different reference gene number choices. (a) Mean accuracy plot of using different reference genes. (b) Break-down run time for different references. Run time is the execution time in this cluster environment and input data: 108 assembled references, 2.4Gb input reads, 64Cores AMD EPYC™ 7702, 512GB RAM.

Another factor that may alter the runtime of the pipeline is the reference panel size. For instance, with large panels (871 species across all angiosperm orders and major families) and extensive gene selections adding to the computational load. However, a balanced setup - using around 100 reference species and a reduced subset of 30 to 50 target genes - enabled the pipeline to run effectively within 5 minutes without significant loss in accuracy (Figure 6a, b), highlighting a feasible compromise between precision and resource demands for general applications.

### Taxonomical specific performance

Among 100 in-silico mixes, prediction accuracy varies in different groups of flowering plants (Figure 7). We provided a summary of performance on selected orders using different numbers of genes as references. We also examined the assembled length for each 353 genes to investigate their lineage specific distribution. The results demonstrate the taxonomical-specific performance of the pipeline, emphasizing its efficiency in different plant lineages. Figure 7a reveals lineage-specific mean accuracy (ACC), showing variations across groups such as Malvales, Lamiales, and Poales. The accuracy, calculated as the average across a range of Z values (0 to 1), indicates some taxonomic groups achieve consistently higher ACC, explaining the better performance in predictions. Meanwhile, we also checked the relative proportion of genes successfully assembled across all 353 genes. Figure 7b shows the recovery proportion for reference genes contains variations in recovery rates, suggesting differences in the performance depending on lineage-specific genetic characteristics and reference gene availability. Taxonomic groups with higher recovery rates exhibit more reliable results, while those with lower rates may require further optimization or additional reference data to enhance pipeline efficiency.

**Figure 7.**
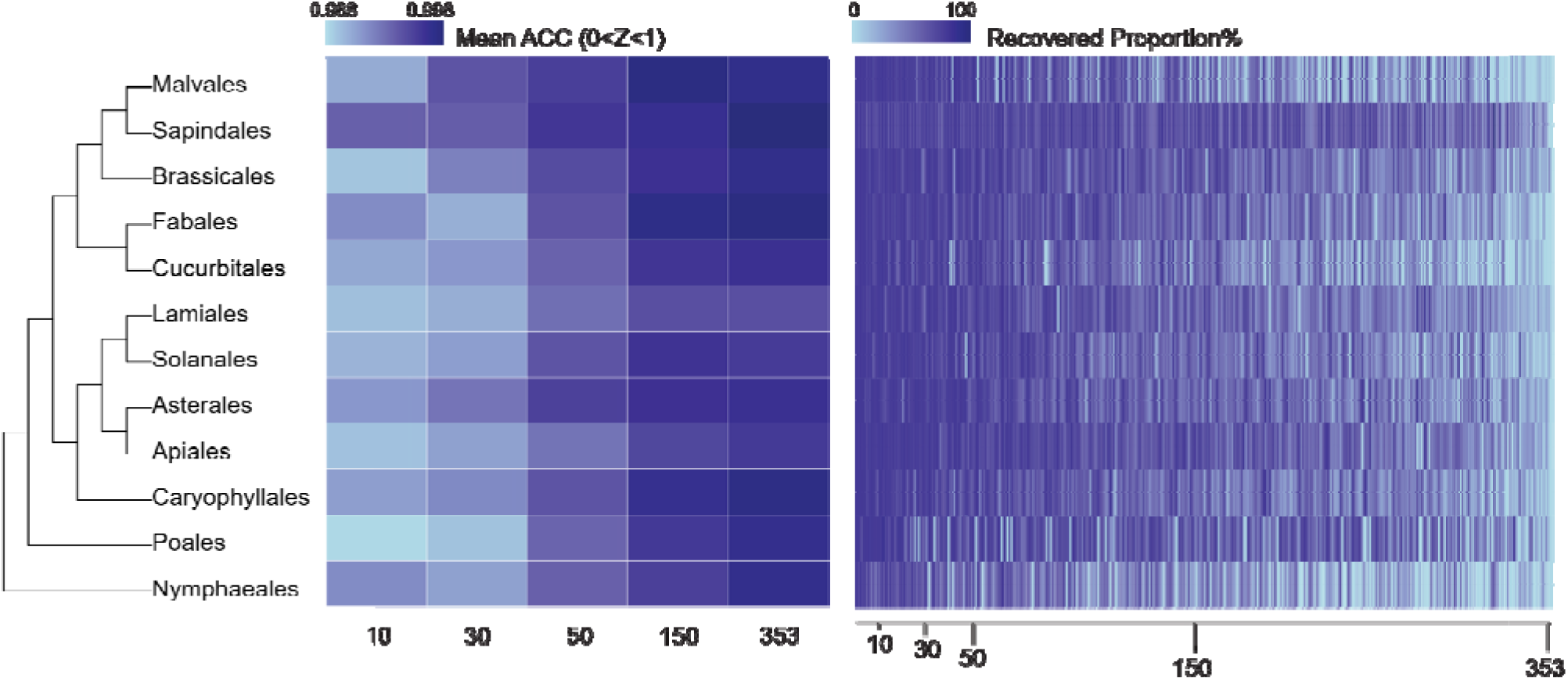
Lineage specific efficiency of the pipeline. (a) Mean ACC in different taxonomy groups. The phylogeny is an ASTRAL tree generated from 30 genes. Blue tiles represent Accuracy value at different reference gene levels. The Accuracy is the averaged number from Z equals to 0 to 1. (b) Heatmap of reference gene mapping rate in different taxonomy groups. Colors represent the percentage of length being recovered. Genes are ordered by occurrence in Angiosperm353 target file.

### Reference gene and species panel design

A comprehensive reference panel is essential for the accurate identification of taxa within mixed samples. Missing lineages in the reference panel can lead to misidentification, as the phylogenetic predictions rely heavily on the diversity and coverage of the reference species included. In specific scenarios, such as detecting contaminants in a particular taxonomic group, designing a targeted reference panel (e.g., focusing on Ericales and Asterales when detecting suspicious contaminants of sunflowers in tea samples; focusing on Poaceae and Fabaceae when detecting potential invasive Mesquite in grassland) can increase computational efficiency and improve precision. Moreover, a hierarchical reference approach offers a practical solution: by first using a broad reference set to identify major clades (e.g., monocots vs. eudicots, commelinid, orders) and subsequently narrowing down with lineage-specific references (families, tribes, genera), the pipeline can reach to an enhanced accuracy and reduced computational demands, especially in high-throughput contexts.

### Technical limitations and potential sources of error

This pipeline has a reasonably high accuracy, but we revealed a few limitations in application. For instance, the sensitivity to read quality and depth, particularly in low-yield scenarios, underscores the importance of optimizing library preparation and sequencing protocols. At low read depths, minor species can drop below detection thresholds, as seen in tests where the mapping rate fell below 20,000 mapped reads. Similarly, the accuracy of exon recovery and tree construction diminishes when read depth is insufficient, potentially impacting ACS calculations and taxonomic predictions. Addressing these limitations may involve adapting pipeline parameters (e.g., gene set size or ACS threshold) or optimizing sequencing strategies for challenging sample types.

Further, the computational complexity of the phylogenetic reconstruction and genetic distance matrix calculations poses challenges, particularly with large, diverse reference sets. Optimizing pipeline efficiency remains an area for improvement, especially as databases like the Kew Gardens Angiosperms353 continue to grow. Expanding the reference panel will undoubtedly improve accuracy across taxonomic groups, but at the cost of increased processing time and computational requirements.

## CONCLUSIONS

### Pipeline Accuracy, Sensitivity, and Practical Implications

This study presents a comprehensive pipeline that combines the Angiosperms353 target sequencing kit with HybPiper for reliable and cost-effective plant species identification in mixed DNA samples. Our results confirm the high -accuracy of Angiosperms353 in both single species and mixed plant DNA identification, shown by both in-silico datasets and real plant samples.

The pipeline achieved overall great evaluating metrices across both in-silico and known mixed samples, highlighting its robustness even in complex mixtures where species proportions varied significantly. Notably, the use of Adjusted Cumulative Similarity (ACS) scores, derived from phylogenetic distance matrices, enabled reliable taxonomic predictions at various hierarchical levels, with sensitivity (TPR), specificity (TNR), precision (PPV) and Accuracy (ACC) optimized around a proper Z-score range. This ACS-based scoring approach is a creative empirical method to assess whether the pipeline can accurately identify closely related species while minimizing false positives and negatives. This is a critical capability for practical applications that demand nuanced taxonomic resolution, such as biodiversity assessments and forensic botany.

### Expanding Applications and Future Directions

This pipeline offers extensive customization possibilities, making it adaptable to a wide range of applications. Modifying gene sets, ACS thresholds, and taxonomic levels provide flexibility in addressing different sample complexities and research needs. For instance, targeted applications such as monitoring agricultural contaminants, regulation of the food supply, detecting contamination and adulteration in foods, dietary supplements, identifying endangered species in soil seed banks, or tracking invasive plant species in mixed environments are well within reach of this workflow. With advancements in reference databases, including de novo sequencing efforts, the scope can be extended to encompass a broader array of plant taxa, enhancing its utility in food science, ecology and conservation biology.

Future directions for this pipeline include exploring machine learning models for pre-filtering candidate species, streamlining algorithms for phylogenetic inference, reducing the need for exhaustive tree construction while maintaining taxonomic resolution, refining the ACS calculation models and expanding the reference database. Incorporating models that consider sequence quality as a variable in the ACS could further refine predictions, especially in low-abundance scenarios. Additionally, a larger reference panel, potentially supplemented by new de novo sequences from projects like OneKP and JGI, will enhance species resolution and identification accuracy across broader clades.

In summary, this study presents a robust, flexible pipeline that addresses the unique challenges of plant species identification in mixed samples. By combining the Angiosperms353 probe set with a streamlined computational approach, this pipeline advances plant metabarcoding as a cost-effective, accurate tool for a variety of scientific and practical applications. Its adaptability and customization potential make it an invaluable asset for studies in conservation biology, environmental monitoring, and food safety, expanding the boundaries of plant species identification in complex sample matrices.

## AUTHOR CONTRIBUTIONS

N.H. implemented software and drafted the manuscript. N.H., M.G.J., and S.H. discussed the initial workflow. M.B. Y.C., and C.M. provided preliminary testing data. C.J. curated reference sequences. C.H. validated the single species workflow Y.C. validated the testing datasets. E.H. and S.H. provided mock supplement mixtures. M.B. generated target sequencing array for real mixes. All authors discussed and approved the final manuscript.

## Acknowledgements

ACKOWLEDGMENTS

This work is funded by FDA – Broad Agency Agreement Contracts Awards (75F40122C00168) under the subject of “Improving detection of plant contaminants in mixed samples with targeted sequencing of 353 nuclear protein coding genes” awarded to M.G.J.

## DATA AVAILABILITY STATEMENT

The pipeline can be found on project GitHub repository (https://github.com/nhu92/SPrOUT). Raw in silico reads were downloaded from ENA (https://www.ebi.ac.uk/ena/browser/home). The taxonomy and phylogenetic information were confirmed through Kew Garden Tree of Life Species Browser (https://treeoflife.kew.org/specimen-viewer). Artificial mixed reads are available on DryAd (DOI: 10.5061/dryad.f4qrfj799). mock supplement mixtures reads were generated from Hunter et al. via Angiosperms353 target sequencing procedure. The mixed component is described in Table S3 and Table S4. The reads for mixed samples are available in NCBI BioProject PRJNA325670. The updated target file and reference for phylogenetic placement is available on project GitHub repository (https://github.com/nhu92/SPrOUT).

## Notes

https://github.com/nhu92/SPrOUT

### Competing Interest Statement

The authors have declared no competing interest.

## References

Alsos, I. G., Y. Lammers, N. G. Yoccoz, T. Jørgensen, P. Sjögren, L. Gielly, and M. E. Edwards. 2018. Plant DNA metabarcoding of lake sediments: How does it represent the contemporary vegetation. Plos One 13: e0195403.

Antil, S., J. S. Abraham, S. Sripoorna, S. Maurya, J. Dagar, S. Makhija, P. Bhagat, et al. 2023. DNA barcoding, an effective tool for species identification: a review. Mol Biol Rep 50: 761–775.

Arstingstall, K. A., S. J. DeBano, X. Li, D. E. Wooster, M. M. Rowland, S. Burrows, and K. Frost. 2021. Capabilities and limitations of using DNA metabarcoding to study plant–pollinator interactions. Molecular Ecology 30: 5266–5297.

Baker, W. J., S. Dodsworth, F. Forest, S. W. Graham, M. G. Johnson, A. McDonnell, L. Pokorny, et al. 2021. Exploring Angiosperms353: An open, community toolkit for collaborative phylogenomic research on flowering plants. Am J Bot 108: 1059–1065.

Bell, K. L., N. d. Vere, A. Keller, R. T. Richardson, A. Gous, K. S. Burgess, and B. J. Brosi. 2016. Pollen DNA barcoding: current applications and future prospects. Genome 59: 629–640.

Bolyen, E., J. R. Rideout, M. R. Dillon, N. A. Bokulich, C. C. Abnet, G. A. Al-Ghalith, H. Alexander, et al. 2019. Reproducible, interactive, scalable and extensible microbiome data science using QIIME 2. Nature Biotechnology 37: 852–857.

Bortolus, A. 2008. Error cascades in the biological sciences: the unwanted consequences of using bad taxonomy in ecology. Ambio 37: 114–118.

Bruno, A., A. Sandionigi, G. Agostinetto, L. Bernabovi, J. Frigerio, M. Casiraghi, and M. Labra. 2019. Food Tracking Perspective: DNA Metabarcoding to Identify Plant Composition in Complex and Processed Food Products. Genes 10: 248.

Bullock, M. R., M. Fokar, R. D. Creek, E. A. Stevens, and M. G. Johnson. 2025. Fun-Sized Library Prep: Miniaturization is a valid method for per-sample cost reduction in targeted sequencing of angiosperm DNA. bioRxiv 10.1101/2025.09.17.676862 DOI: 2025.2009.2017.676862.

Camacho, C., G. Coulouris, V. Avagyan, N. Ma, J. Papadopoulos, K. Bealer, and T. L. Madden. 2009. BLAST+: architecture and applications. BMC Bioinformatics 10: 421.

Capella-Gutierrez, S., J. M. Silla-Martinez, and T. Gabaldon. 2009. trimAl: a tool for automated alignment trimming in large-scale phylogenetic analyses. Bioinformatics 25: 1972–1973.

Chac, L. D., and B. B. Thinh. 2023. Species Identification through DNA Barcoding and Its Applications: A Review. Biology Bulletin 50: 1143–1156.

Chen, S. 2023. Ultrafast one-pass FASTQ data preprocessing, quality control, and deduplication using fastp. iMeta 2: e107.

Crisci, J. V., L. Katinas, M. J. Apodaca, and P. C. Hoch. 2020. The End of Botany. Trends Plant Sci 25: 1173–1176.

Fazekas, A. J., K. S. Burgess, P. R. Kesanakurti, S. W. Graham, S. G. Newmaster, B. C. Husband, D. M. Percy, M. Hajibabaei, and S. C. H. Barrett. 2008. Multiple Multilocus DNA Barcodes from the Plastid Genome Discriminate Plant Species Equally Well. Plos One 3: e2802.

Goodstein, D. M., S. Shu, R. Howson, R. Neupane, R. D. Hayes, J. Fazo, T. Mitros, et al. 2012. Phytozome: a comparative platform for green plant genomics. Nucleic Acids Res 40: D1178–1186.

Group1, C. P. B., D.-Z. Li, L.-M. Gao, H.-T. Li, H. Wang, X.-J. Ge, J.-Q. Liu, et al. 2011. Comparative analysis of a large dataset indicates that internal transcribed spacer (ITS) should be incorporated into the core barcode for seed plants. Proceedings of the National Academy of Sciences 108: 19641-19646.

Hale, H., E. M. Gardner, J. Viruel, L. Pokorny, and M. G. Johnson. 2020. Strategies for reducing per-sample costs in target capture sequencing for phylogenomics and population genomics in plants. Appl Plant Sci 8: e11337.

Hebert, P. D. N., R. Floyd, S. Jafarpour, and S. W. J. Prosser. 2025. Barcode 100K Specimens: In a Single Nanopore Run. Molecular Ecology Resources 25: e14028.

Hollingsworth, P. M., S. W. Graham, and D. P. Little. 2011. Choosing and Using a Plant DNA Barcode. Plos One 6: e19254.

Hunter, E. S., R. Literman, and S. M. Handy. 2021. Utilizing Big Data to Identify Tiny Toxic Components: Digitalis. Foods 10: 1794.

Johnson, M. G., E. M. Gardner, Y. Liu, R. Medina, B. Goffinet, A. J. Shaw, N. J. Zerega, and N. J. Wickett. 2016. HybPiper: Extracting coding sequence and introns for phylogenetics from high-throughput sequencing reads using target enrichment. Appl Plant Sci 4: 1600016.

Johnson, M. G., L. Pokorny, S. Dodsworth, L. R. Botigue, R. S. Cowan, A. Devault, W. L. Eiserhardt, et al. 2019. A Universal Probe Set for Targeted Sequencing of 353 Nuclear Genes from Any Flowering Plant Designed Using k-Medoids Clustering. Syst Biol 68: 594–606.

Kalyaanamoorthy, S., B. Q. Minh, T. K. F. Wong, A. von Haeseler, and L. S. Jermiin. 2017. ModelFinder: fast model selection for accurate phylogenetic estimates. Nature Methods 14: 587–589.

Katoh, K., and M. C. Frith. 2012. Adding unaligned sequences into an existing alignment using MAFFT and LAST. Bioinformatics 28: 3144–3146.

Lee, A. K., I. S. Gilman, M. Srivastav, A. D. Lerner, M. J. Donoghue, and W. L. Clement. 2021. Reconstructing Dipsacales phylogeny using Angiosperms353: issues and insights. Am J Bot 108: 1122–1142.

Li, H., B. Handsaker, A. Wysoker, T. Fennell, J. Ruan, N. Homer, G. Marth, et al. 2009. The Sequence Alignment/Map format and SAMtools. Bioinformatics 25: 2078–2079.

Liu, M., L. J. Clarke, S. C. Baker, G. J. Jordan, and C. P. Burridge. 2020. A practical guide to DNA metabarcoding for entomological ecologists. Ecological Entomology 45: 373–385.

Maurin, O., A. Anest, S. Bellot, E. Biffin, G. Brewer, T. Charles-Dominique, R. S. Cowan, et al. 2021. A nuclear phylogenomic study of the angiosperm order Myrtales, exploring the potential and limitations of the universal Angiosperms353 probe set. Am J Bot 108: 1087–1111.

Minh, B. Q., H. A. Schmidt, O. Chernomor, D. Schrempf, M. D. Woodhams, A. von Haeseler, and R. Lanfear. 2020. IQ-TREE 2: New Models and Efficient Methods for Phylogenetic Inference in the Genomic Era. Mol Biol Evol 37: 1530–1534.

Newbold, T. 2018. Future effects of climate and land-use change on terrestrial vertebrate community diversity under different scenarios. Proc Biol Sci 285: 20180792.

Norgaard, L., C. R. Olesen, K. Trojelsgaard, C. Pertoldi, J. L. Nielsen, P. Taberlet, A. Ruiz-Gonzalez, M. De Barba, and L. Iacolina. 2021. eDNA metabarcoding for biodiversity assessment, generalist predators as sampling assistants. Sci Rep 11: 6820.

One Thousand Plant Transcriptomes, I. 2019. One thousand plant transcriptomes and the phylogenomics of green plants. Nature 574: 679–685.

Perez-Escobar, O. A., S. Dodsworth, D. Bogarin, S. Bellot, J. A. Balbuena, R. J. Schley, I. A. Kikuchi, et al. 2021. Hundreds of nuclear and plastid loci yield novel insights into orchid relationships. Am J Bot 108: 1166–1180.

Pezzini, F. F., G. Ferrari, L. L. Forrest, M. L. Hart, K. Nishii, and C. A. Kidner. 2023. Target capture and genome skimming for plant diversity studies. Applications in Plant Sciences 11: e11537.

Piper, A. M., J. Batovska, N. O. I. Cogan, J. Weiss, J. P. Cunningham, B. C. Rodoni, and M. J. Blacket. 2019. Prospects and challenges of implementing DNA metabarcoding for high-throughput insect surveillance. Gigascience 8.

Price, M. N., P. S. Dehal, and A. P. Arkin. 2009. FastTree: computing large minimum evolution trees with profiles instead of a distance matrix. Mol Biol Evol 26: 1641–1650.

Schloss, P. D., S. L. Westcott, T. Ryabin, J. R. Hall, M. Hartmann, E. B. Hollister, R. A. Lesniewski, et al. 2009. Introducing mothur: Open-Source, Platform-Independent, Community-Supported Software for Describing and Comparing Microbial Communities. Applied and Environmental Microbiology 75: 7537–7541.

Schutte, A., P. E. Stuben, and J. J. Astrin. 2023. Molecular Weevil Identification Project: A thoroughly curated barcode release of 1300 Western Palearctic weevil species (Coleoptera, Curculionoidea). Biodivers Data J 11: e96438 (eng).

Shen, W., B. Sipos, and L. Zhao. 2024. SeqKit2: A Swiss army knife for sequence and alignment processing. iMeta 3: e191.

Siniscalchi, C. M., O. Hidalgo, L. Palazzesi, J. Pellicer, L. Pokorny, O. Maurin, I. J. Leitch, et al. 2021. Lineage-specific vs. universal: A comparison of the Compositae1061 and Angiosperms353 enrichment panels in the sunflower family. Appl Plant Sci 9.

Slater, G. S. C., and E. Birney. 2005. Automated generation of heuristics for biological sequence comparison, BMC Bioinformatics, 31.

Slimp, M., L. D. Williams, H. Hale, and M. G. Johnson. 2021. On the potential of Angiosperms353 for population genomic studies. Appl Plant Sci 9.

Thomas, A. E., J. Igea, H. M. Meudt, D. C. Albach, W. G. Lee, and A. J. Tanentzap. 2021a. Using target sequence capture to improve the phylogenetic resolution of a rapid radiation in New Zealand Veronica. Am J Bot 108: 1289–1306.

Thomas, S. K., X. Liu, Z. Y. Du, Y. Dong, A. Cummings, L. Pokorny, Q. J. Xiang, and J. H. Leebens-Mack. 2021b. Comprehending Cornales: phylogenetic reconstruction of the order using the Angiosperms353 probe set. Am J Bot 108: 1112–1121.

Vallin, H., H. Hipperson, J. Titěra, L. Jones, and M. Fraser. 2025. Comparative Analysis of Pasture Composition: DNA Metabarcoding Versus Quadrat-Based Botanical Surveys in Experimental Grassland Plots. Ecology and Evolution 15: e71195.

Wizenberg, S. B., L. R. Newburn, M. Pepinelli, I. M. Conflitti, R. T. Richardson, S. E. R. Hoover, R. W. Currie, P. Giovenazzo, and A. Zayed. 2023. Validating a multi-locus metabarcoding approach for characterizing mixed-pollen samples. Plant Methods 19: 120.

Yu, X.-Q., Y.-Z. Jiang, R. A. Folk, J.-L. Zhao, C.-N. Fu, L. Fang, H. Peng, J.-B. Yang, and S.-X. Yang. 2022. Species discrimination in Schima (Theaceae): Next-generation super-barcodes meet evolutionary complexity. Molecular Ecology Resources 22: 3161–3175.

Zimmer, E. A., and J. Wen. 2015. Using nuclear gene data for plant phylogenetics: Progress and prospects II. Next-gen approaches. Journal of Systematics and Evolution 53: 371–379.

Zuntini, A. R., L. P. Frankel, L. Pokorny, F. Forest, and W. J. Baker. 2021. A comprehensive phylogenomic study of the monocot order Commelinales, with a new classification of Commelinaceae. Am J Bot 108: 1066–1086.

Zuntini, A. R., T. Carruthers, O. Maurin, P. C. Bailey, K. Leempoel, G. E. Brewer, N. Epitawalage, et al. 2024. Phylogenomics and the rise of the angiosperms. Nature 629: 843–850.

