## Supplemental Tables for "SPrOUT: A computational and targeted sequencing approach for mixed plant DNA identification with Angiosperms353"

Table S1. Testing species datasets for 100 in silico mixes.

| ID | Scientific Name | ERR Series | Reads Number | Recovered Gene Number | Recovered Base Pair | Order | Family |
|---|---|---|---|---|---|---|---|
| 1 | Brasenia schreberi | ERR7618329 | 1500860 | 325 | 146967 | Nymphaeales | Nymphaeaceae |
| 2 | Celosia trigyna | ERR7621855 | 5015608 | 342 | 187602 | Caryophyllales | Amaranthaceae |
| 3 | Turbinicarpus valdezianus | ERR7619841 | 7603762 | 344 | 191130 | Caryophyllales | Cactaceae |
| 4 | Polycarpaea repens | ERR7621975 | 4102970 | 345 | 170847 | Caryophyllales | Caryophyllaceae |
| 5 | Ribes rubrum | ERR7621129 | 5838268 | 349 | 203742 | Saxifragales | Grossulariaceae |
| 6 | Darmera peltata | ERR7621478 | 3414216 | 350 | 187395 | Saxifragales | Saxifragaceae |
| 7 | Alepidea amatymbica | ERR5034284 | 2429578 | 343 | 171960 | Apiales | Apiaceae |
| 8 | Tetrapanax papyrifer | ERR7619972 | 5039516 | 348 | 207354 | Apiales | Araliaceae |
| 9 | Aphanactis jamesoniana | ERR6041621 | 1644776 | 343 | 189462 | Asterales | Asteraceae |
| 10 | Galactites tomentosus | ERR7621201 | 7462824 | 345 | 185796 | Asterales | Asteraceae |
| 11 | Tarchonanthus trilobus | ERR7621921 | 3695578 | 349 | 209937 | Asterales | Asteraceae |
| 12 | Colquhounia coccinea | ERR9230289 | 4620680 | 350 | 231732 | Lamiales | Lamiaceae |
| 13 | Hesperelaea palmeri | ERR10978383 | 1799488 | 350 | 215976 | Lamiales | Oleaceae |
| 14 | Bonamia spectabilis | ERR7621887 | 3039352 | 345 | 179112 | Solanales | Convolvulaceae |
| 15 | Eriolarynx australis | ERR7622024 | 1846154 | 343 | 161928 | Solanales | Solanaceae |
| 16 | Chesneya yunnanensis | ERR7618004 | 1533038 | 340 | 188454 | Fabales | Fabaceae |
| 17 | Securidaca diversifolia | ERR7621813 | 4536730 | 346 | 181152 | Fabales | Polygalaceae |
| 18 | Corallocarpus welwitschii | ERR7619688 | 6576798 | 349 | 184845 | Cucurbitales | Cucurbitaceae |
| 19 | Gynostemma pentaphyllum | ERR7619700 | 5763930 | 343 | 174453 | Cucurbitales | Cucurbitaceae |
| 20 | Gironniera subaequalis | ERR7621631 | 5188400 | 347 | 207636 | Rosales | Cannabaceae |
| 21 | Pouzolzia rubricaulis | ERR5034939 | 1993008 | 341 | 163401 | Rosales | Urticaceae |
| 22 | Firmiana simplex | ERR7622322 | 3125336 | 350 | 189585 | Malvales | Malvaceae |
| 23 | Englerodaphne subcordata | ERR5034905 | 3300628 | 336 | 158493 | Malvales | Thymelaeaceae |
| 24 | Dithyrea californica | SRR22519649 | 4736586 | 347 | 261942 | Brassicales | Brassicaceae |
| 25 | Neocalyptrocalyx muco | ERR7622178 | 5264540 | 348 | 213111 | Brassicales | Cleomaceae |
| 26 | Citrus hystrix | ERR7620065 | 5183344 | 350 | 216060 | Sapindales | Rutaceae |
| 27 | Caladium lindenii | ERR7618821 | 3460438 | 325 | 122208 | Alismatales | Araceae |
| 28 | Agropyropsis lolium | ERR7621428 | 1213096 | 326 | 153225 | Poales | Poaceae |
| 29 | Dinochloa orenuda | ERR10978135 | 2542272 | 342 | 228243 | Poales | Poaceae |
| 30 | Heteranthoecia guineensis | ERR7619905 | 6287468 | 344 | 215874 | Poales | Poaceae |

Table S2. Species composition of 100 in silico mixes.

| Individual IDs | Mixed Species Number |
|---|---|
| 12x18x3 | 3 |
| 12x1x15 | 3 |
| 12x25x2 | 3 |
| 12x25x4 | 3 |
| 12x7x10 | 3 |
| 13x10x1 | 3 |
| 13x24x8 | 3 |
| 13x28x15 | 3 |
| 13x7x10 | 3 |
| 15x10x28 | 3 |
| 15x13x28 | 3 |
| 15x22x12 | 3 |
| 15x8x28 | 3 |
| 16x13x1 | 3 |
| 16x28x8 | 3 |
| 18x10x22 | 3 |
| 18x25x12 | 3 |
| 18x7x13 | 3 |
| 18x9x28 | 3 |
| 19x22x10 | 3 |
| 19x28x4 | 3 |
| 1x2x13 | 3 |
| 1x30x25 | 3 |
| 1x4x19 | 3 |
| 22x26x4 | 3 |
| 24x28x1 | 3 |
| 24x4x28 | 3 |
| 24x9x15 | 3 |
| 26x19x8 | 3 |
| 26x1x24 | 3 |
| 28x16x30 | 3 |
| 2x15x24 | 3 |
| 2x22x19 | 3 |
| 2x24x15 | 3 |
| 30x4x3 | 3 |
| 3x7x13 | 3 |
| 4x15x10 | 3 |
| 7x26x30 | 3 |
| 8x7x1 | 3 |
| 8x7x3 | 3 |
| 10x19x28x9x15x22 | 6 |
| 10x22x30x16x13x26 | 6 |
| 10x24x7x19x2x25 | 6 |
| 10x26x9x24x13x2 | 6 |

| | |
|---|---|
| 10x9x15x8x25x18 | 6 |
| 12x13x1x18x22x7 | 6 |
| 12x22x15x26x3x1 | 6 |
| 12x25x22x2x26x10 | 6 |
| 18x19x28x30x7x12 | 6 |
| 19x25x4x26x2x1 | 6 |
| 19x7x8x16x3x10 | 6 |
| 1x18x8x28x26x2 | 6 |
| 1x7x16x18x25x8 | 6 |
| 24x10x28x1x25x13 | 6 |
| 25x12x28x2x10x13 | 6 |
| 25x1x19x28x10x24 | 6 |
| 25x28x2x15x3x8 | 6 |
| 25x30x15x19x3x13 | 6 |
| 26x3x2x1x4x10 | 6 |
| 28x12x25x10x3x15 | 6 |
| 28x19x2x9x18x26 | 6 |
| 28x19x4x2x13x3 | 6 |
| 28x24x25x15x4x1 | 6 |
| 2x1x28x15x3x7 | 6 |
| 2x22x25x7x24x3 | 6 |
| 30x10x8x4x25x26 | 6 |
| 30x4x13x26x19x28 | 6 |
| 30x4x16x1x22x3 | 6 |
| 30x4x19x7x3x16 | 6 |
| 30x9x22x28x16x12 | 6 |
| 3x16x28x24x1x7 | 6 |
| 4x19x30x2x13x16 | 6 |
| 4x28x12x16x3x19 | 6 |
| 4x7x8x15x16x1 | 6 |
| 7x10x4x16x13x15 | 6 |
| 7x28x25x10x4x30 | 6 |
| 8x16x15x7x3x9 | 6 |
| 8x16x9x25x3x18 | 6 |
| 8x30x16x22x15x19 | 6 |
| 9x19x12x16x7x25 | 6 |
| 10x4x13x19x2x15x1x9 | 10 |
| 12x1x7x4x3x2x13x18x | 10 |
| 12x9x19x18x26x25x1 | 10 |
| 13x26x15x9x28x8x10x | 10 |
| 16x25x10x24x12x8x30 | 10 |
| 1x19x8x22x25x30x7x2 | 10 |
| 1x25x24x22x28x19x1 | 10 |
| 25x16x10x1x12x9x18x | 10 |
| 25x28x30x7x22x24x1 | 10 |
| 26x22x3x13x10x19x4x | 10 |
| 2x12x28x10x1x9x30x3 | 10 |

| | |
|---|---|
| 2x13x24x3x12x7x16x1 | 10 |
| 2x24x25x9x28x16x4x7 | 10 |
| 30x28x8x2x9x25x15x4 | 10 |
| 4x28x15x7x16x9x13x1 | 10 |
| 4x30x25x16x10x3x28x | 10 |
| 4x3x16x25x18x22x2x8 | 10 |
| 7x22x30x8x18x13x3x1 | 10 |
| 9x15x18x2x25x8x10x1 | 10 |
| 9x30x1x4x18x13x22x8 | 10 |

Table S3. Parameter sets as the preparation for testing low yield samples (Poales).

| Raw Input Reads Number | Random Seed | Targeted Mapped Reads Number |
|---|---|---|
| 344827 | 100 | 100k |
| 344827 | 100 | 100k |
| 344827 | 200 | 100k |
| 344827 | 200 | 100k |
| 344827 | 300 | 100k |
| 344827 | 300 | 100k |
| 172413 | 100 | 50k |
| 172413 | 100 | 50k |
| 172413 | 200 | 50k |
| 172413 | 200 | 50k |
| 172413 | 300 | 50k |
| 172413 | 300 | 50k |
| 86206 | 100 | 25k |
| 86206 | 100 | 25k |
| 86206 | 200 | 25k |
| 86206 | 200 | 25k |
| 86206 | 300 | 25k |
| 86206 | 300 | 25k |
| 68965 | 100 | 20k |
| 68965 | 100 | 20k |
| 68965 | 200 | 20k |
| 68965 | 200 | 20k |
| 68965 | 300 | 20k |
| 68965 | 300 | 20k |
| 51724 | 100 | 15k |
| 51724 | 100 | 15k |
| 51724 | 200 | 15k |
| 51724 | 200 | 15k |
| 51724 | 300 | 15k |
| 51724 | 300 | 15k |
| 34483 | 100 | 10k |
| 34483 | 100 | 10k |
| 34483 | 200 | 10k |
| 34483 | 200 | 10k |
| 34483 | 300 | 10k |
| 34483 | 300 | 10k |

Table S4. Parameter sets as the preparation for testing low yield samples with low proportions (Asterales).

| Raw Input Reads Number | Random Seed | Targeted Mapped Reads Number | Targeted Mapped Reads Proportion (%) |
|---|---|---|---|
| 2777750 | 100 | 100k | 10 |
| 2777750 | 100 | 100k | 10 |
| 5555500 | 100 | 100k | 5 |
| 5555500 | 100 | 100k | 5 |
| 11111000 | 100 | 100k | 2.5 |
| 11111000 | 100 | 100k | 2.5 |
| 1388861 | 100 | 50k | 10 |
| 1388861 | 100 | 50k | 10 |
| 2777722 | 100 | 50k | 5 |
| 2777722 | 100 | 50k | 5 |
| 5555444 | 100 | 50k | 2.5 |
| 5555444 | 100 | 50k | 2.5 |
| 694416 | 100 | 25k | 10 |
| 694416 | 100 | 25k | 10 |
| 1388833 | 100 | 25k | 5 |
| 1388833 | 100 | 25k | 5 |
| 2777666 | 100 | 25k | 2.5 |
| 2777666 | 100 | 25k | 2.5 |
| 555527 | 100 | 20k | 10 |
| 555527 | 100 | 20k | 10 |
| 1111055 | 100 | 20k | 5 |
| 1111055 | 100 | 20k | 5 |
| 2222111 | 100 | 20k | 2.5 |
| 2222111 | 100 | 20k | 2.5 |
| 416638 | 100 | 15k | 10 |
| 416638 | 100 | 15k | 10 |
| 833277 | 100 | 15k | 5 |
| 833277 | 100 | 15k | 5 |
| 1666555 | 100 | 15k | 2.5 |
| 1666555 | 100 | 15k | 2.5 |
| 277777 | 100 | 10k | 10 |
| 277777 | 100 | 10k | 10 |
| 555555 | 100 | 10k | 5 |
| 555555 | 100 | 10k | 5 |
| 1111111 | 100 | 10k | 2.5 |
| 1111111 | 100 | 10k | 2.5 |

Table S4. Performance of each species in the 100 mixes test

| Species | Z Score | TPR Mean | TPR SD | TNR Mean | TNR SD | PPV Mean | PPV SD | ACC Mean | ACC SD |
|---|---|---|---|---|---|---|---|---|---|
| 1 | 0.3 | 0.989 | 0.042 | 1 | 0 | 1 | 0 | 0.999 | 0.004 |
| 1 | 0.6 | 0.928 | 0.107 | 1 | 0 | 1 | 0 | 0.993 | 0.011 |
| 1 | 1 | 0.825 | 0.139 | 1 | 0 | 1 | 0 | 0.982 | 0.016 |
| 2 | 0.3 | 0.994 | 0.031 | 0.996 | 0.013 | 0.97 | 0.096 | 0.996 | 0.012 |
| 2 | 0.6 | 0.975 | 0.063 | 1 | 0 | 1 | 0 | 0.998 | 0.006 |
| 2 | 1 | 0.88 | 0.135 | 1 | 0 | 1 | 0 | 0.987 | 0.015 |
| 3 | 0.3 | 0.994 | 0.031 | 0.994 | 0.015 | 0.954 | 0.128 | 0.994 | 0.015 |
| 3 | 0.6 | 0.97 | 0.081 | 0.999 | 0.007 | 0.986 | 0.074 | 0.996 | 0.01 |
| 3 | 1 | 0.85 | 0.148 | 1 | 0 | 1 | 0 | 0.984 | 0.016 |
| 4 | 0.3 | 1 | 0 | 0.997 | 0.011 | 0.979 | 0.085 | 0.997 | 0.01 |
| 4 | 0.6 | 0.97 | 0.066 | 1 | 0 | 1 | 0 | 0.997 | 0.007 |
| 4 | 1 | 0.854 | 0.144 | 1 | 0 | 1 | 0 | 0.985 | 0.017 |
| 7 | 0.3 | 0.995 | 0.03 | 0.994 | 0.017 | 0.944 | 0.154 | 0.993 | 0.016 |
| 7 | 0.6 | 0.967 | 0.082 | 0.997 | 0.012 | 0.963 | 0.12 | 0.993 | 0.014 |
| 7 | 1 | 0.863 | 0.142 | 0.999 | 0.006 | 0.987 | 0.072 | 0.984 | 0.016 |
| 8 | 0.3 | 0.994 | 0.031 | 0.997 | 0.01 | 0.968 | 0.097 | 0.996 | 0.01 |
| 8 | 0.6 | 0.981 | 0.049 | 0.999 | 0.003 | 0.991 | 0.047 | 0.997 | 0.007 |
| 8 | 1 | 0.859 | 0.147 | 1 | 0 | 1 | 0 | 0.984 | 0.019 |
| 9 | 0.3 | 1 | 0 | 0.999 | 0.004 | 0.992 | 0.036 | 0.999 | 0.004 |
| 9 | 0.6 | 0.983 | 0.045 | 1 | 0 | 1 | 0 | 0.998 | 0.006 |
| 9 | 1 | 0.862 | 0.157 | 1 | 0 | 1 | 0 | 0.982 | 0.021 |
| 10 | 0.3 | 1 | 0 | 0.998 | 0.01 | 0.98 | 0.09 | 0.998 | 0.009 |
| 10 | 0.6 | 0.976 | 0.061 | 0.999 | 0.003 | 0.993 | 0.043 | 0.997 | 0.008 |
| 10 | 1 | 0.867 | 0.156 | 1 | 0 | 1 | 0 | 0.984 | 0.021 |
| 12 | 0.3 | 1 | 0 | 1 | 0 | 1 | 0 | 1 | 0 |
| 12 | 0.6 | 0.984 | 0.056 | 1 | 0 | 1 | 0 | 0.998 | 0.007 |
| 12 | 1 | 0.912 | 0.128 | 1 | 0 | 1 | 0 | 0.989 | 0.016 |
| 13 | 0.3 | 1 | 0 | 0.988 | 0.02 | 0.885 | 0.188 | 0.988 | 0.019 |
| 13 | 0.6 | 0.988 | 0.038 | 0.995 | 0.012 | 0.941 | 0.133 | 0.994 | 0.012 |
| 13 | 1 | 0.885 | 0.147 | 0.999 | 0.006 | 0.988 | 0.071 | 0.985 | 0.02 |
| 15 | 0.3 | 1 | 0 | 0.997 | 0.009 | 0.97 | 0.102 | 0.997 | 0.009 |
| 15 | 0.6 | 0.968 | 0.07 | 0.999 | 0.004 | 0.984 | 0.062 | 0.995 | 0.009 |
| 15 | 1 | 0.889 | 0.155 | 1 | 0 | 1 | 0 | 0.986 | 0.02 |
| 16 | 0.3 | 0.994 | 0.032 | 0.999 | 0.005 | 0.978 | 0.079 | 0.998 | 0.005 |
| 16 | 0.6 | 0.975 | 0.077 | 1 | 0 | 1 | 0 | 0.997 | 0.008 |
| 16 | 1 | 0.823 | 0.155 | 1 | 0 | 1 | 0 | 0.981 | 0.019 |
| 18 | 0.3 | 0.993 | 0.035 | 0.996 | 0.013 | 0.967 | 0.114 | 0.996 | 0.013 |
| 18 | 0.6 | 0.988 | 0.041 | 0.998 | 0.011 | 0.978 | 0.104 | 0.996 | 0.011 |
| 18 | 1 | 0.885 | 0.13 | 0.998 | 0.007 | 0.983 | 0.083 | 0.985 | 0.018 |
| 19 | 0.3 | 1 | 0 | 0.996 | 0.012 | 0.972 | 0.093 | 0.996 | 0.011 |
| 19 | 0.6 | 0.959 | 0.077 | 1 | 0 | 1 | 0 | 0.995 | 0.009 |
| 19 | 1 | 0.843 | 0.148 | 1 | 0 | 1 | 0 | 0.984 | 0.017 |
| 22 | 0.3 | 1 | 0 | 0.998 | 0.008 | 0.989 | 0.053 | 0.998 | 0.007 |
| 22 | 0.6 | 0.979 | 0.057 | 1 | 0 | 1 | 0 | 0.997 | 0.008 |
| 22 | 1 | 0.901 | 0.13 | 1 | 0 | 1 | 0 | 0.988 | 0.017 |
| 24 | 0.3 | 0.992 | 0.036 | 0.998 | 0.008 | 0.974 | 0.091 | 0.997 | 0.008 |

| | | | | | | | | | |
|---|---|---|---|---|---|---|---|---|---|
| 24 | 0.6 | 0.936 | 0.115 | 0.999 | 0.004 | 0.989 | 0.053 | 0.993 | 0.011 |
| 24 | 1 | 0.819 | 0.159 | 1 | 0 | 1 | 0 | 0.979 | 0.021 |
| 25 | 0.3 | 1 | 0 | 0.999 | 0.003 | 0.996 | 0.025 | 0.999 | 0.003 |
| 25 | 0.6 | 0.97 | 0.071 | 1 | 0 | 1 | 0 | 0.997 | 0.008 |
| 25 | 1 | 0.883 | 0.134 | 1 | 0 | 1 | 0 | 0.987 | 0.016 |
| 26 | 0.3 | 0.991 | 0.039 | 0.999 | 0.004 | 0.991 | 0.039 | 0.998 | 0.005 |
| 26 | 0.6 | 0.973 | 0.064 | 1 | 0 | 1 | 0 | 0.997 | 0.006 |
| 26 | 1 | 0.923 | 0.14 | 1 | 0 | 1 | 0 | 0.99 | 0.02 |
| 28 | 0.3 | 0.991 | 0.039 | 0.996 | 0.012 | 0.95 | 0.129 | 0.995 | 0.012 |
| 28 | 0.6 | 0.941 | 0.103 | 0.999 | 0.004 | 0.986 | 0.058 | 0.993 | 0.01 |
| 28 | 1 | 0.847 | 0.163 | 1 | 0 | 1 | 0 | 0.984 | 0.018 |
| 30 | 0.3 | 1 | 0 | 0.997 | 0.009 | 0.971 | 0.085 | 0.997 | 0.008 |
| 30 | 0.6 | 0.986 | 0.04 | 1 | 0 | 1 | 0 | 0.998 | 0.006 |
| 30 | 1 | 0.898 | 0.126 | 1 | 0 | 1 | 0 | 0.987 | 0.017 |

Table S6. Species composition of 60 in silico mixes for family level test.

| Within Order Mixes | Cross 2 Orders Mixes | Cross 3 Orders Mixes |
|---|---|---|
| 02x03 | 02x09 | 01x09x28 |
| 02x04 | 02x10 | 01x09x29 |
| 03x04 | 02x11 | 01x09x30 |
| 09x10 | 03x09 | 01x10x28 |
| 09x11 | 03x10 | 01x10x29 |
| 10x11 | 03x11 | 01x10x30 |
| 28x29 | 04x09 | 01x11x28 |
| 28x30 | 04x10 | 01x11x29 |
| 29x30 | 04x11 | 01x11x30 |
| 02x03x04 | 02x28 | 02x09x28 |
| 09x10x11 | 02x29 | 02x09x29 |
| 28x29x30 | 02x30 | 02x09x30 |
| | 03x28 | 02x10x28 |
| | 03x29 | 02x10x29 |
| | 03x30 | 02x10x30 |
| | 04x28 | 02x11x28 |
| | 04x29 | 02x11x29 |
| | 04x30 | 02x11x30 |
| | 09x28 | 03x09x28 |
| | 09x29 | 03x09x29 |
| | 09x30 | 03x09x30 |
| | 10x28 | 03x10x28 |
| | 10x29 | 03x10x29 |
| | 10x30 | 03x10x30 |
| | 11x28 | 03x11x28 |
| | 11x29 | 03x11x29 |
| | 11x30 | 03x11x30 |

Table S7. Real mixes predicting results in order and family levels.

| Mix Name | Library ID | Component # | Species | Weight (g) | Tissue ID | Tissue Type | Proportion | Expected Orders | Expected Families | Predicted Orders | Predicted Families |
|---|---|---|---|---|---|---|---|---|---|---|---|
| brassica | EM-1 | 1 | Brassica juncea | 0.0154 | NYBG 02757110 | leaf | 12.39936 | Brassicales | Brassicaceae | Brassicales | Brassicaceae |
| brassica | EM-1 | 2 | Brassica napus var. pabularia | 0.0155 | wen12944 | leaf | 12.47987 | Brassicales | Brassicaceae | Brassicales | Brassicaceae |
| brassica | EM-1 | 3 | Brassica nigra | 0.016 | NYBG 01282366 | leaf | 12.88245 | Brassicales | Brassicaceae | Brassicales | Brassicaceae |
| brassica | EM-1 | 4 | Brassica oleracea | 0.0155 | wen12943 | leaf | 12.47987 | Brassicales | Brassicaceae | Brassicales | Brassicaceae |
| brassica | EM-1 | 5 | Brassica oleracea var. capitata | 0.0156 | wen17939 | leaf | 12.56039 | Brassicales | Brassicaceae | Brassicales | Brassicaceae |
| brassica | EM-1 | 6 | Brassica rapa subsp. chinensis | 0.015 | wen17943 | leaf | 12.07729 | Brassicales | Brassicaceae | Brassicales | Brassicaceae |
| brassica | EM-1 | 7 | Brassica rapa subsp. pekinensis | 0.0157 | wen17942 | leaf | 12.6409 | Brassicales | Brassicaceae | Brassicales | Brassicaceae |
| brassica | EM-1 | 8 | Brassica rapa subsp. rapa | 0.0155 | wen17977 | leaf | 12.47987 | Brassicales | Brassicaceae | Brassicales | Brassicaceae |
| prunus | EM-2 | 1 | Prunus armeniaca | 0.0153 | wen13871 | leaf | 12.37864 | Rosales | Rosaceae | Rosales | Rosaceae |
| prunus | EM-2 | 2 | Prunus avium | 0.0158 | wen13831 | leaf | 12.78317 | Rosales | Rosaceae | Rosales | Rosaceae |
| prunus | EM-2 | 3 | Prunus davidiana | 0.0156 | wen13690 | leaf | 12.62136 | Rosales | Rosaceae | Rosales | Rosaceae |
| prunus | EM-2 | 4 | Prunus dulcis | 0.0157 | wen13509 | leaf | 12.70227 | Rosales | Rosaceae | Rosales | Rosaceae |
| prunus | EM-2 | 5 | Prunus mume | 0.0152 | wen14010 | leaf | 12.29773 | Rosales | Rosaceae | Rosales | Rosaceae |
| prunus | EM-2 | 6 | Prunus padus | 0.016 | wen17760 | leaf | 12.94498 | Rosales | Rosaceae | Rosales | Rosaceae |
| prunus | EM-2 | 7 | Prunus persica | 0.0155 | wen13868 | leaf | 12.54045 | Rosales | Rosaceae | Rosales | Rosaceae |
| prunus | EM-2 | 8 | Prunus yedoensis | 0.0145 | NYBG 02759465 | leaf | 11.73139 | Rosales | Rosaceae | Rosales | Rosaceae |
| citrus | EM-3 | 1 | Citrus maxima | 0.0163 | wen13444 | leaf | 19.70979 | Sapindales | Rutaceae | Sapindales | Rutaceae |
| citrus | EM-3 | 2 | Citrus medica | 0.0168 | wen12888 | leaf | 20.31439 | Sapindales | Rutaceae | Sapindales | Rutaceae |
| citrus | EM-3 | 3 | Citrus reticulata | 0.0155 | NCNPR 20739 | leaf | 18.74244 | Sapindales | Rutaceae | Sapindales | Rutaceae |
| citrus | EM-3 | 4 | Citrus sinensis | 0.0341 | AHP Lot #3729F | fruit | 41.23337 | Sapindales | Rutaceae | Sapindales | Rutaceae |
| vaccinium | EM-4 | 1 | Vaccinium corymbosum | 0.0158 | wen12931 | leaf | 33.19328 | Ericales | Ericaceae | Ericales | Ericaceae |
| vaccinium | EM-4 | 2 | Vaccinium darrowii | 0.0164 | NYBG 01426190 | leaf | 34.45378 | Ericales | Ericaceae | Ericales | Ericaceae |
| vaccinium | EM-4 | 3 | Vaccinium macrocarpon | 0.0154 | wen14928a | leaf | 32.35294 | Ericales | Ericaceae | Ericales | Ericaceae |
| mix1 | EM-5 | 1 | Carthamus tinctorius | 0.0163 | AHP Lot #4879 | leaf | 10.75908 | Asterales | Asteraceae | . | . |
| mix1 | EM-5 | 2 | Beta vulgaris | 0.0153 | AHP | leaf | 10.09901 | Caryophyllales | Amararthaceae | Caryophyllales | Amararthaceae |
| mix1 | EM-5 | 3 | Curcuma longa | 0.0155 | wen13706 | leaf | 10.23102 | Zingiberales | Zingiberaceae | Zingiberales | Zingiberaceae |
| mix1 | EM-5 | 4 | Daucus carota subsp. sativus | 0.0161 | wen13408 | leaf | 10.62706 | Apiales | Apiaceae | Apiales | Apiaceae |
| mix1 | EM-5 | 5 | Zea mays | 0.0161 | wen17880 | leaf | 10.62706 | Poales | Poaceae | Poales | Poaceae |
| mix1 | EM-5 | 6 | Gardenia jasminoides | 0.0159 | wen17593 | leaf | 10.49505 | Gentianales | Rubiaceae | Gentianales | Rubiaceae |
| mix1 | EM-5 | 7 | Nelumbo nucifera | 0.0563 | AHP | root | 37.16172 | Proteales | Nelumbonaceae | . | . |
| mix2 | EM-6 | 1 | Scutellaria baicalensis | 0.0161 | GF-15 | leaf | 20.85492 | Lamiales | Lamiaceae | Lamiales | Lamiaceae |
| mix2 | EM-6 | 2 | Panax ginseng | 0.0153 | wen3116 | leaf | 19.81865 | Apiales | Araliaceae | Apiales | Araliaceae |
| mix2 | EM-6 | 3 | Zingiber officinale | 0.0152 | wen13758 | leaf | 19.68912 | Zingiberales | Zingiberaceae | Zingiberales | Zingiberaceae |
| mix2 | EM-6 | 4 | Camellia sinensis | 0.0151 | wen13341 | leaf | 19.55959 | Ericales | Theaceae | . | . (No reference) |
| mix2 | EM-6 | 5 | Ocimum tenuiflorum | 0.0155 | AHP Lot # 5818 | leaf | 20.07772 | Lamiales | Lamiaceae | Lamiales | Lamiaceae |
| mix3 | EM-7 | 1 | Ginkgo biloba | 0.0253 | wen13439 | leaf | 49.7053 | Ginkgoales | Ginkgoaceae | . (No reference) | . (No reference) |
| mix3 | EM-7 | 2 | Zizania latifolia | 0.0256 | wen Tibet 923 | leaf | 50.2947 | Poales | Poaceae | Poales | Poaceae |
| mix5 | EM-8 | 1 | Artemisia annua | 0.0159 | GF-5 | leaf | 24.76636 | Asterales | Asteraceae | Asterales | Asteraceae |
| mix5 | EM-8 | 2 | Coptis chinensis | 0.0158 | wen12843 | leaf | 24.61059 | Ranunculales | Ranunculaceae | Ranunculales | Ranunculaceae |
| mix5 | EM-8 | 3 | Centella asiatica | 0.0166 | Jim Duke 6 | leaf | 25.8567 | Apiales | Apiaceae | Apiales | Apiaceae |
| mix5 | EM-8 | 4 | Gynostemma pentaphyllum | 0.0159 | wen13272 | leaf | 24.76636 | Cucurbitales | Cucurbitaceae | Cucurbitales | Cucurbitaceae |
| mix6 | EM-9 | 1 | Litchi chinensis | 0.0157 | wen13292 | leaf | 16.66667 | Sapindales | Sapindaceae | Sapindales | Sapindaceae |
| mix6 | EM-9 | 2 | Dimocarpus longan | 0.0153 | wen13297 | leaf | 16.24204 | Sapindales | Sapindaceae | Sapindales | Sapindaceae |
| mix6 | EM-9 | 3 | Citrus maxima | 0.0154 | wen13444 | leaf | 16.3482 | Sapindales | Rutaceae | Sapindales | Rutaceae |
| mix6 | EM-9 | 4 | Citrus reticulata | 0.0163 | NCNPR 20736 | leaf | 17.30361 | Sapindales | Rutaceae | Sapindales | Rutaceae |
| mix6 | EM-9 | 5 | Punica granatum | 0.016 | wen17924 | leaf | 16.98514 | Myrtales | Lythraceae | Myrtales | Lythraceae |
| mix6 | EM-9 | 6 | Psidium guajava | 0.0155 | wen17850 | leaf | 16.45435 | Myrtales | Myrtaceae | Myrtales | Myrtaceae |
| mix7 | EM-10 | 1 | Allium sativum | 0.0153 | Jim Duke 14 | leaf | 24.79741 | Asparagales | Asparagaceae | Asparagales | Amaryllidaceae |
| mix7 | EM-10 | 2 | Asparagus officinalis | 0.015 | wen13722 | leaf | 24.31118 | Asparagales | Asparagaceae | Asparagales | . |
| mix7 | EM-10 | 3 | Brassica oleracea | 0.0153 | wen12943 | leaf | 24.79741 | Brassicales | Brassicaceae | Brassicales | Brassicaceae |
| mix7 | EM-10 | 4 | Artocarpus heterophyllus | 0.0161 | wen13606 | leaf | 26.094 | Rosales | Moraceae | Rosales | Moraceae |
| mix8 | EM-11 | 1 | Theobroma cacao | 0.0156 | wen12890 | leaf | 21.63662 | Malvales | Malvaceae | Malvales | Malvaceae |
| mix8 | EM-11 | 2 | Coffea arabica | 0.0446 | wen12944 | leaf | 61.85853 | Gentianales | Rubiaceae | Gentianales | Rubiaceae |
| mix8 | EM-11 | 3 | Vanilla planifolia | 0.0119 | wen12897 | leaf | 16.50485 | Asparagales | Orchidaceae | . | . (No reference) |
