## Supplemental Figures for "SPrOUT: A computational and targeted sequencing approach for mixed plant DNA identification with Angiosperms353"

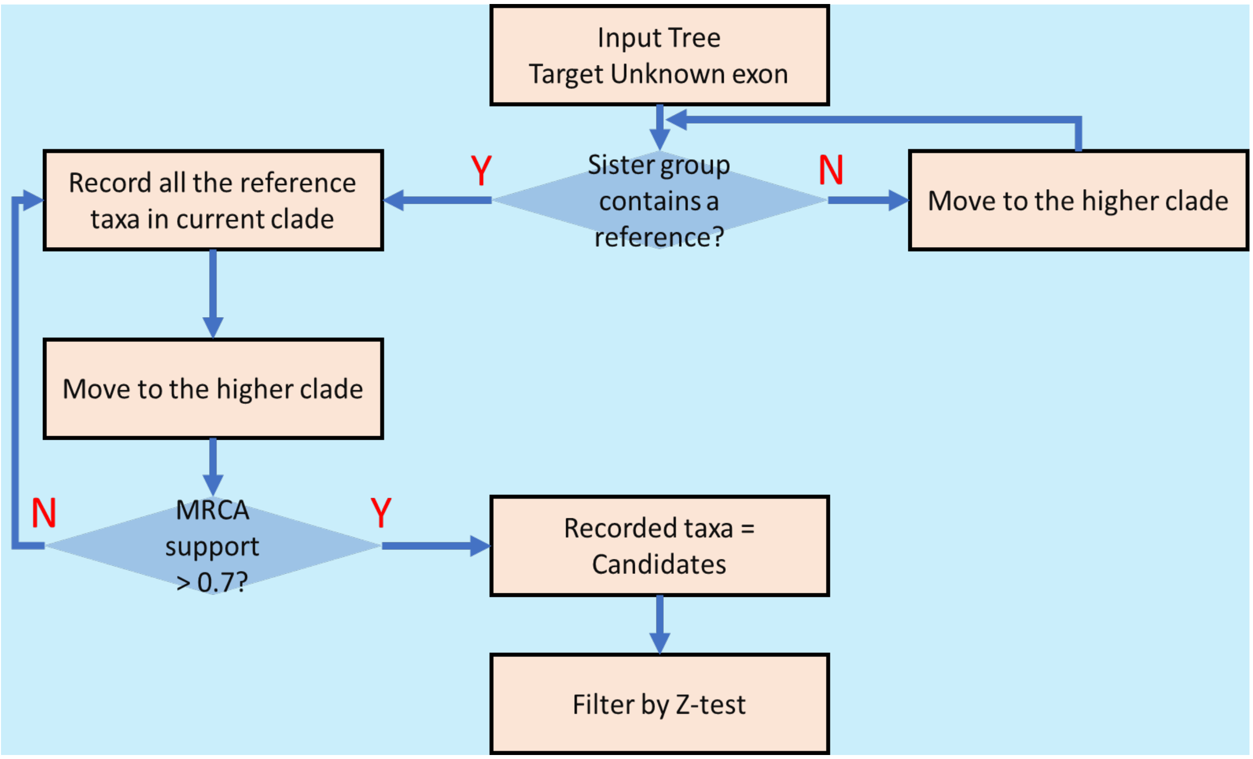


Figure S1. Flowchart of filtering non-related species before final ACS calculation.

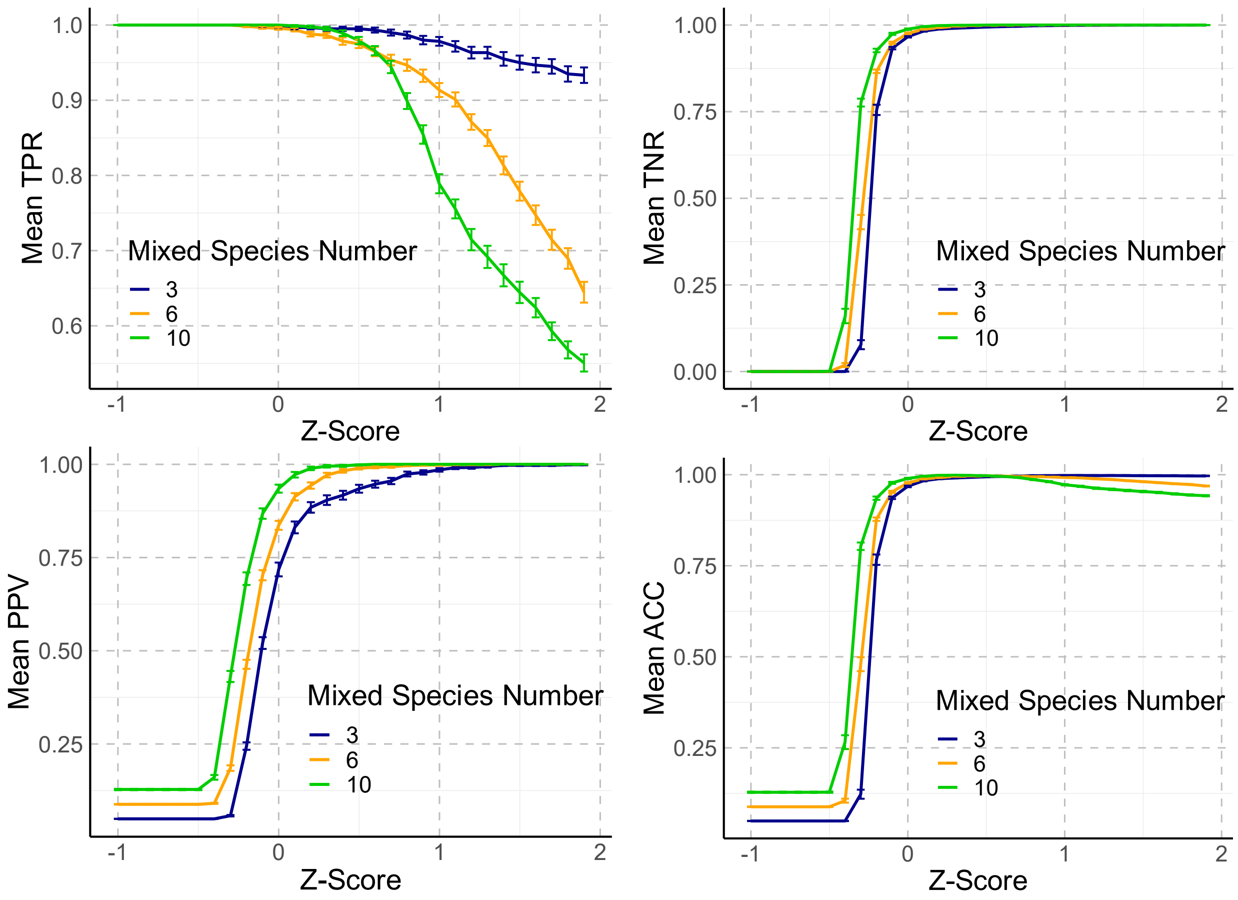


Figure S2. Evaluations of the pipeline using four parameters from the confusion matrix (See Table 1 for Glossaries). TPR = True Position Rate; TNR = True Negative Rate; PPV = Positive Predictive Value; ACC = Accuracy. Colored lines indicate performance for different numbers of mixed species in testing data. Error bar is the standard error.

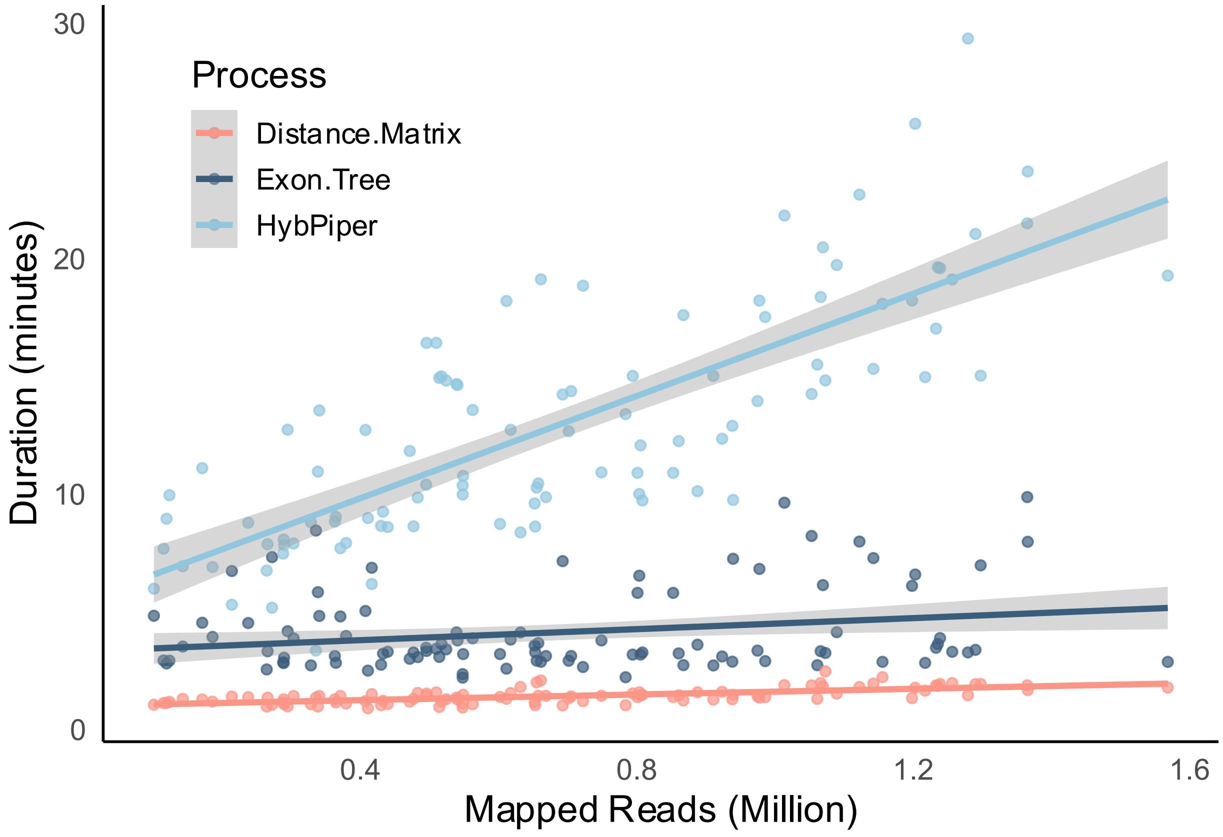


Figure S3. Relationship between mapped read number and processing duration for three steps in the bioinformatic pipeline. Run time is the execution time in this cluster environment and input data: 50 targeted genes, 108 assembled references, 64Cores AMD EPYC™ 7702, 512GB RAM.
